# S100A11 promotes focal adhesion disassembly via myosin II-driven contractility and Piezo1-mediated Ca^2+^ entry

**DOI:** 10.1101/2023.07.17.549432

**Authors:** Tareg Omer Mohammed, You-Rong Lin, Kai Weissenbruch, Kien Xuan Ngo, Yanjun Zhang, Noriyuki Kodera, Martin Bastmeyer, Yusuke Miyanari, Azuma Taoka, Clemens M. Franz

**Affiliations:** WPI Nano Life Science Institute, Kanazawa University, Kanazawa, Japan; Cell and Neurobiology, Zoological Institute, Karlsruhe Institute of Technology, Karlsruhe, Germany; Institute for Biological and Chemical Systems – Biological Information Processing, Karlsruhe Institute of Technology, Eggenstein-Leopoldshafen, Germany; Cancer Research Institute, Kanazawa University, Kanazawa, Japan; Institute of Science and Engineering, Kanazawa University, Kanazawa, Japan

**Keywords:** S100A11, focal adhesion, actin stress fiber, myosin II, Piezo1, Ca^2+^

## Abstract

S100A11 is a small Ca^2+^-activatable protein with an established role in different cellular processes involving actin cytoskeleton remodeling, such as cell migration, membrane protrusion formation, and plasma membrane repair. It also displays Ca^2+^-dependent F-actin binding activity and localizes to actin stress fibers (SFs), but its precise role in regulating these structures remains unclear. Analyzing endogenous S100A11 localization in HeLa and U2OS osteosarcoma cells confirmed SF association but in addition revealed steady localization to stable focal adhesions (FAs), typically at the end of dorsal stress fibers. In contrast, S100A11 levels at FAs increased sharply, but transiently, at the onset of peripheral FA disassembly. Elevating intracellular Ca^2+^ levels using the Ca^2+^ ionophore ionomycin reliably stimulated both S100A11 recruitment and subsequent FA disassembly. However, pre-incubation with the non-muscle myosin II (NM II) inhibitor blebbistatin, or with an inhibitor to the stretch-activatable Ca^2+^ channel Piezo1 effectively suppressed S100A11 recruitment, implicating S100A11 in an actomyosin contractility-driven FA disassembly mechanism involving Piezo1-dependent Ca^2+^ influx. Applying external mechanical forces on peripheral FAs via a micropipette likewise recruited S100A11 to FAs, even when NM II was inhibited by blebbistatin or in NM IIA knockout cells, corroborating the mechanosensitive recruitment mechanism of S100A11. However, extracellular Ca^2+^ and Piezo1 function was still indispensable, indicating that NM II-dependent contraction forces act upstream of Piezo1-mediated Ca^2+^ influx, in turn leading to S100A11 activation and FA recruitment. Moreover, S100A11 knockout cells feature enlarged FAs and display delayed FA disassembly during cell membrane retraction, consistent with impaired FA turnover in these cells. Our results thus demonstrate a novel mechano-sensitive function for S100A11 in promoting actomyosin contractility-driven FA disassembly.

## Introduction

S100 proteins form a family of small calcium-binding proteins with multifaceted roles in diverse intra- and extracellular processes, including calcium homeostasis, signal transduction, proliferation, differentiation, and migration (Donato et al., 2013; Gonzalez et al., 2020; Moore, 1965), as well as inflammation (Sreejit et al., 2020) and cancer (Bresnick et al., 2015). Central to the role of S100 proteins as signaling molecules is their capacity to become activated by Ca^2+^ binding (Schäfer and Heizmann, 1996). S100 proteins typically form symmetric homodimers (Yap et al., 1999), in which each subunit supplies two central EF-hand motifs connected by a short hinge region. Binding of Ca^2+^ via the EF hands induces a conformational change that exposes a normally inaccessible hydrophobic cleft for interaction with other target proteins (Otterbein et al., 2002; Santamaria-Kisiel et al., 2006). Ca^2+^ greatly increases the affinity of S100 proteins to their targets, and consequently S100 proteins have been described as intracellular Ca^2+^ sensors, able to detect and translate fluctuations in intracellular Ca^2+^ levels into specific cellular responses (Zimmer and Weber, 2010). In addition to the common EF hand motifs arrangement, individual S100 members contain short C-terminal tails that vary in length and sequence and give each member additional specific functional roles.

S100A11, also called S100C or calgizzarin (Ohta et al., 1991; Todoroki et al., 1991), is widely expressed and like other S100 family members plays a role in numerous cellular functions under both physiological and pathological conditions (He et al., 2009). Amongst other functions, it is associated with cell cycle and cell growth regulation, and apoptosis (Sakaguchi and Huh, 2011; Sakaguchi et al., 2003; Xia et al., 2018). S100A11 also has a well-established role in a variety of diseases, including cancer metastasis, inflammation, and neurological diseases (Zhang et al., 2021). S100A11 binds filamentous actin (F-actin) in a Ca^2+^-dependent manner (Sakaguchi et al., 2000; Zhao et al., 2000) and participates in different cellular processes involving actin cytoskeleton remodeling, including membrane wound healing (Jaiswal et al., 2014), pseudopodal protrusion formation (Shankar et al., 2010), and cell migration (Meng et al., 2019; Wang et al., 2013). Furthermore, S100A11 localizes along actin stress fibers (SFs), filopodia, and lamellipodia in normal and immortalized human fibroblasts, implicating it in actin organization and regulation of dynamics (Sakaguchi et al., 2000). S100A11 also inhibits smooth muscle myosin ATPase activity in a Ca^2+^-dependent manner (Zhao et al., 2000), pointing towards a potential role in regulating actomyosin contractility. Furthermore, Ca^2+^ entry into migrating cancer cells as a result of plasma membrane injury recruits S100A11 together with annexin A2, another Ca^2+^-binding protein, to the site of injury, where the S100A11/annexin A2 complexes help resealing the membrane by stimulating cortical actin polymerization (Jaiswal et al., 2014). Similarly, S100A11 and annexin A2 also cooperate during actin-dependent wound healing in endothelial cells (Ashraf and Gerke, 2021). Complexes between S100A11 and annexin 6 on the other hand may stabilize connections between the plasma membrane and the cytoskeleton (Chang et al., 2007). S100A11 also drives the formation of actin-dependent pseudopodal protrusions in migrating tumor cells (Shankar et al., 2010).

Despite the ample evidence demonstrating S100A11/F-actin interaction and S100A11-dependent actin cytoskeleton regulation, previous research has focused more strongly on other S100 members, particularly S100A4, S100A6, and S100A10 in relation to actin cytoskeleton dynamics (Gómez-Contreras et al., 2017; Jurewicz et al., 2020; Sayeed et al., 2013). As a result, little is known about the molecular mechanisms by which S100A11 modulates actin remodeling. Here we have investigated the intracellular localization of S100A11 in HeLa and U2OS osteosarcoma cells and detected a novel localization to stable peripheral focal adhesions (FAs). Moreover, S100A11 levels at FAs increase sharply, yet transiently, at the onset of FA disassembly, implicating S100A11 in FA turnover regulation. Recruitment of S100A11 to disassembling FAs requires NM II-mediated contractility, opening of stretch-activatable Piezo1 Ca^2+^ channels at stressed FAs, and activation by intracellular Ca^2+^. Furthermore, we demonstrate aberrant FA morphology and impaired FA disassembly in S100A11 knockout (KO) cells. Together, our results thus reveal a novel mechano-regulated role for S100A11 during NM II-, Piezo1-, and Ca^2+^-dependent FA disassembly.

## Materials and Method

### Cell culture and plasmid transfection

Human osteosarcoma U2OS cells kindly provided by Tsukasa Matsunaga, Kanazawa University, HeLa cells were obtained from the ATCC. Cells were cultured in Dulbecco’s modified Eagle’s medium (DMEM) supplemented with 10% fetal bovine serum (FBS), 1% L-glutamine, penicillin (10,000 U/ml) and streptomycin sulfate (10,000 µg/ml). For maintenance, cells were passaged two times a week and incubated in a humidified incubator at 37°C, 5% CO_2_. Transient plasmid transfection was performed using Lipofectamine 3000 (ThermoFisher) according to the manufacturer’s instructions. Lifeact-mCherry was a gift from Moritoshi Sato (Addgene plasmid #67302 (Kawano et al., 2015)), GFP-S100A11 was a gift from Volker Gerke and Ursula Rescher (Addgene plasmid #107201), and Piezo1-GFP was a gift from Satoshi Arai (Addgene plasmid #80925 (Coste et al., 2010)). The vinculin-mCherry construct has been published previously (Carisey et al., 2013) and R-GECO1 was a kind gift from Satoshi Arai (Addgene plasmid #32444 (Zhao et al., 2011a)).

### Cloning and protein purification

To produce recombinant His-tag GFP-S100A11, cDNA encoding S100A11 from a human cDNA library was amplified by PCR using the forward primer 5’-ACTATACGGTGGATCCatggcaaaaatctccagcc-3’ and the reverse primer 5’-GCAGGTCGACAAGCTTtcaggtccgcttct-3’ purchased from Macrogen, Japan. The N-terminal His-tag (x6His) GFP-S100A11 construct was then generated by infusion cloning. S100A11 was inserted into the linear PcoldI His-tag GFP plasmid, which was cut using BamHI and HindIII sites. The nucleotide sequence of the PcoldI His-tag GFP-S100A11 expression vector was confirmed by DNA sequencing and restriction enzyme digestion. Protein expression was performed in pG-KJE8 E. coli BL21 (Takara, Japan) using the protocol provided by the manufacturer. Purification of the His-tag GFP-S100A11 protein was performed as described in a previous study (Ngo et al., 2015).

### CRISPR/Cas9 S100A11 knockout

The U2OS S100A11 KO cell line was generated by genome editing using a pCas9P plasmid expressing Cas9 and two sgRNA sequences (#57891 & #57892, both on exon2 of S100A11) selected from the Brunello human CRISPR knockout library (Doench et al., 2016). The sgRNAsequences were: 5′ - AGAACTAGCTGCCTTCACAA - 3′, and 5′ CCA GCC CTA CAG AGA CTGAG - 3′. After cloning of the S100A11-KO plasmid, 1 µg of the constructed plasmid was transfected into U2OS cells using Lipofectamin 3000 Transfection Reagent. The following day, the transfection medium was removed, and cells were cultured in growth medium containing 10 µg/mL puromycin for three days to select for knockout cells. Finally, single clones were isolated by limited dilution into 96 well plates and then expanded. Absence of S100A11 expression was confirmed by Western Blotting using Rabbit anti-S100A11 (ProteinTech Group).

### Immunofluorescence

For immunofluorescence staining cells were cultured in uncoated 35 mm glass bottom dishes (either FD35 Fluorodish, WPI, or 1130H, Matsunami Glass) for 24 h to 48 h and then fixed in 4% paraformaldehyde solution in PBS for 10 min at RT. This was followed by simultaneous blocking and permeabilization in a solution composed of 50 mM glycine, 0.05% Tween 20, 0.1% Triton X-100 and 0.1% BSA in PBS for 30 min. Primary antibodies (Rabbit Anti-S100A11, GeneTex, Mouse monoclonal anti-β-tubulin, Sigma-Aldrich, clone TUB 2.1, and mouse anti-Vinculin Merck, V9131) were then diluted in PBS containing 10 mM glycine, 0.05% Tween 20, 0.1% Triton X-100, and cells were incubated for 2h at RT or overnight at 4 ℃. After three washes with PBS to remove unbound primary antibodies, cells were stained with the appropriate secondary antibodies (goat anti-rabbit AF568, goat anti-rabbit AF488, goat anti-mouse AF568, or goat anti-mouse AF488, all Invitrogen) diluted 200× in 0.1% Tween 20 in PBS for 30 min. Phalloidin AF568 or Phalloidin AF488 (Invitrogen) diluted at 1:200 were added along with secondary antibodies when cells were co-stained for F-actin. Finally, cells were washed three times with PBS and imaged using the 60x objective of an inverted fluorescence phase contrast microscope (BZ-X810 Keyence All-in-one Fluorescence Microscope).

### TIRF live cell imaging

Total internal reflection fluorescence (TIRF) microscopy was performed using a Nikon Ti-E inverted microscope equipped with a 100× CFI Apo TIRF objective lens and 488 nm and 561 nm lasers (Sapphire, Coherent). A high-sensitivity electron-multiplying charge-coupled-device camera (iXon3; Andor, DU897E-CS0) with electron multiplying and preamplifier gains of 296 and 2.4×, respectively, was used in combination with a 1.5× C-mount adapter (Nikon) to acquire images. The Z-position was adjusted for optimal focus and maintained using the Nikon Perfect Focus System during time-lapse imaging. Cells were imaged at 28°C under ambient air conditions in culture medium containing 20 mM HEPES (pH 7.4). Timelapse series were stabilized, and histograms were equalized in Fiji. Kymographs were generated using the “KymoResliceWide” or “Multi Kymograph” Fiji plugins. Fluorescence intensity plots were generated in Fiji after correcting for photobleaching.

### Western blotting

Cell lysates were prepared by washing cells once with ice-cold PBS and then scrapping the cells in 200 μl Triton X-100 lysis buffer containing 1% (vol/vol) Triton X-100, 150 mM NaCl, 50 mM Tris-HCl pH 7.4 and Complete Protease Inhibitor Cocktail (Thermo Scientific). The mixture was kept for 30 min on ice and then centrifuged at 12,000 rpm for 20 min at 4°C. 20 μl of protein sample was mixed with 4 μl of 6x SDS sample buffer, boiled at 95°C for 5 min, and then loaded onto 10– 20% gradient Mini-PROTEAN TGX Precast gels (Bio-Rad) for SDS-PAGE. The electrophoresis was first run for ∼40 min at 80 V, followed by 150 V for ∼ 60 min. After electrophoresis, proteins were transferred onto 0.2 µm PVDF membranes in a wet tank apparatus (Bio-Rad) run at 100 V for 1 h at room temperature while cooling with an ice pack in a Tris-Glycine transfer buffer containing 20 mM Tris, 200 mM glycine, and 20% (v/v) methanol. Subsequently, the membrane was blocked with 5% non-fat dried milk in TBS-T (20 mM Tris-HCl pH 7.4, 150 mM NaCl, 0.05% (v/v) Tween 20) for 1 h and then incubated with primary antibodies (Rabbit Anti-S100A11 (ProteinTech), and Mouse monoclonal TUB 2.1 anti-β-Tubulin (Sigma-Aldrich) at 1000× dilution in blocking buffer overnight at 4°C. After 3 washes in TBS-T for 10 min, blots were incubated with HRP-conjugated secondary antibodies (Goat anti-Rabbit HRP (Invitrogen), or Goat anti-Mouse HRP (SouthernBiotech) at 2000× dilution in blocking buffer for 1 h. After 3 final washes in TBS-T, blots were developed using the Western Lightning Plus-ECL chemiluminescence substrate and imaged using an AE-9300 Ez capture MG (ATTO, Tokyo, Japan) imaging system.

### Inhibitor and drug treatments

For ionomycin-induced FA disassembly, cells were grown on glass bottom dishes for 16h. Cells were then transferred to the TIRF microscopy platform and 3 µM ionomycin (Merck) was added directly during live-cell image acquisition. Typically, cells were imaged for 1 h until FA disassembly had completed. To assess the effect of NM IIA inhibition on ionomycin-induced GFPSA11 accumulation at FA, HeLa cells transiently transfected with S100A11-GFP were transferred onto the TIRF stage and imaged for 30 min in the presence of 50 µM blebbistatin (Merck). After addition of ionomycin (final concentration 3 µM), cells were imaged for a further 20 ∼ 30 min. The effect of NM IIA inhibition on ionomycin induced GFPSA11 accumulation was then assessed by comparing the fluorescence intensity of blebbistatin treated cells with that of untreated control cells (5 cells per group). The effect of NM IIA inhibition on cell contraction was measured by treating untransfected U2OS cells with 50 µM of blebbistatin followed by immediate live-cell imaging by holographic tomography (3D Explorer, NanoLive). After 30 min of imaging in blebbistatin-containing medium, cells were treated with 3 µM ionomycin and imaged for an additional 20 ∼ 30 min. To examine the role of Piezo1 in mediating Ca^2+^ influx and S100A11 recruitment near stressed FAs, GFP-S100A11 transfected cells were treated with 3 μM of the Piezo1 inhibitor GsMTx4 (Peptide Institute) for 3 h. Subsequently, cells were stimulated with 3 μM ionomycin, and the dynamics of GFP-S100A11 was investigated by live cell TIRF imaging. In other experiments, GsMTx4-treated cells were subjected to external force application using the micropipette assay.

### Focal adhesion dynamics analysis

U2OS wildtype and S100A11 KO cells were transfected with vinculin-mCherry using Lipofectamine 3000 (Thermo Fisher). After 24 h, cells were replated in 35-mm glass-bottom dishes coated with 20 μg/mL fibronectin for imaging. Time-lapse image sets were acquired at 1 frame per minute for a total of 60 minutes at 37°C in a chamber providing 5% CO_2_ and an automated fluorescence microscope (KEYENCE BZ-X810). The acquired images were analyzed in ImageJ using background subtraction, frame stabilization, and photobleaching correction plug-ins. To determine FA dynamics, peripheral cell regions displaying active membrane retraction (net cell edge movement >5 µm over 1 h) were identified. Within these areas, 15 x 15 µ*mm*^2^ subregions were selected, which typically contained 10 to 25 FAs. In these regions, FAs were then tracked using the Cellpose TrackMate module in ImageJ (Stringer et al., 2021). Individual FA area and retraction speeds were obtained from at least 70 FAs from 5 different cells per cell type. Non-dynamic FA parameters (individual FA size, FA area per cell) were determined from immunofluorescence images of fixed U2OS wildtype, S100A11 KO, and S100A11 KO/GFP-S100A11 rescue cells by adopting a described method (Horzum et al., 2014). Statistical tests for significance were performed in the GraphPad Prism software or Origin 2021.

### Micropipette pulling force experiments

Micropipette external force application experiments were performed based on a previous report (Riveline et al., 2001). Briefly, glass micropipettes were fabricated by pulling borosilicate glass capillaries (inner diameter 0.58 mm, outer diameter 1.00 mm) into nanopipettes with a tip diameter of ∼100 nm using a CO_2_ laser puller (Model P-2000, Sutter Instruments Co., USA). For external force application experiments, U2OS cells transiently expressing GFP-S100A11 were plated into 35 mm glass bottom dish coated with 10 μg/ml human plasma fibronectin and grown overnight. After transferring cells into growth medium containing 20 mM HEPES, pH 7.4, the cell sample was placed onto an inverted Olympus CKX53 light microscope equipped with a 40x lens (LUCPLFLN40X), a camera (U3-3880CP-M-GL Rev.2.2, IDS) and a U-LGPS fluorescence light source (Olympus). A micropipette mounted on a manual *x*,*y*,*z*-micromanipulator system (Narishige, Japan) was then positioned over a target cell, while the intracellular localization of GFP-S100A11 was recorded by timelapse fluorescence microscopy. The micropipette was then lowered until contacting the plasma membrane of the recorded cell, and subsequently gently pushed against the protruding cell nucleus, thereby exerting pulling forces against peripheral FAs. GFP-S100A11 localization to stressed FA was typically observed within 30 to 180 sec of force application. In some cases, cells were co-transfected with GFP-S100A11 and vinculin-mCherry to verify FA localization. Requirement for non-muscle NM IIA (NM IIA) was further investigated by using U2OS NM IIA knockout cells (Weißenbruch et al., 2021) or by preincubating U2OS wildtype cells with blebbistatin (50 μM) for 30 min before force application. Additional experiments were performed in Ca^2+^-free medium to investigate the requirement of Ca^2+^ for S100A11 translocation.

## Results

### S100A11 localizes to Focal adhesions and stress fibers in different cell types

S100A11 is an actin binding protein (Zhao et al., 2000) previously reported to localize to different elements of the actin cytoskeleton, including filopodia, lamellipodia, and SFs in normal and transformed fibroblast (Sakaguchi et al., 2000). While these localization patterns suggest a general role for S100A11 in actin cytoskeleton regulation, its precise function in these processes remains poorly understood. To further investigate potential roles of S100A11 in cytoskeleton regulation, we analyzed the intracellular localization of endogenous S100A11 in HeLa cells by immunofluorescence (IF). Co-staining for F-actin with phalloidin confirmed the previously reported localization of S100A11 along actin SFs (Fig. 1A). However, in a subset of cells S100A11 was also strongly enriched at SF termini within the cell periphery. Co-staining for vinculin confirmed prominent localization of S100A11 to FAs (Fig. 1B). This observation revealed a previously unknown additional localization of S100A11 to FAs. S100A11 has also been implicated in microtubule-binding (Broome and Eckert, 2004), but co-staining for S100A11 and β-tubulin showed that S100A11 does not localize to microtubules in these cells (Supp. Fig. S1). Nevertheless, the non-overlapping localization patterns of S100A11 and microtubules ruled out potential fluorescence channel cross talk in our setup. In the IF experiments we used an enhanced staining protocol (Flores-Maldonado et al., 2020), which increased the contrast of the obtained S100A11 images compared to standard IF protocols. S100A11 localization to FAs could also be independently confirmed by total internal reflection fluorescence (TIRF) microscopy of transfected HeLa cells expressing GFP-S100A11 (Suppl. Fig. S2A). Since the TIRF imaging z-range is restricted to ∼200 nm above the cell culture substrate, GFP-S100A11 was not detected in SFs, which typically extend higher into the cytoplasmic space, in these experiments. However, both SF and FA localization could easily be detected by epi-fluorescence imaging of GFP-S100A11 expressing cells, even against strong cytoplasmic GFP-S100A11 levels (Supp. Fig. S2B).

**Fig. 1.**
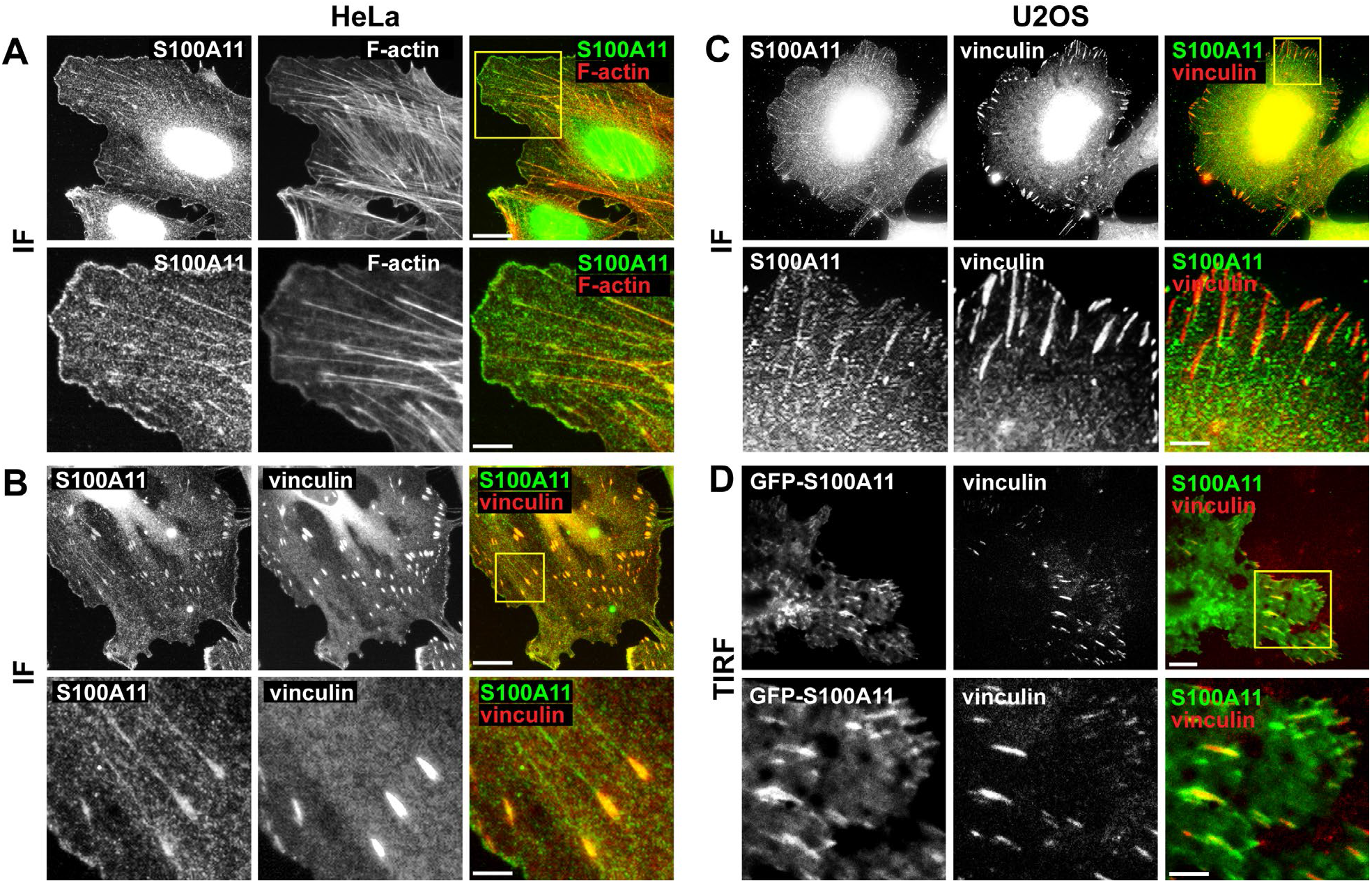
S100A11 localizes to actin stress fibers and focal adhesions. **(A)** Immunostaining for S100A11 and phalloidin staining of F-actin in HeLa cells. Full size images (top row) and cropped images corresponding to the yellow box (bottom row). **(B)** Dual immunostaining for S100A11 and vinculin in HeLa cells. Full size images (top row) and cropped images corresponding to the yellow box (bottom row). **(C)** Dual immunostaining for S100A11 and vinculin in U2OS cells. Full size images (top row) and cropped images corresponding to the yellow box (bottom row). **(D)** TIRF microscopy live-cell images of U2OS cells expressing GFP-S100A11 and vinculin-mCherry. Full size images (top row) and cropped images corresponding to the yellow box (bottom row). Scale bars 10 µm and 2.5 µm (**A, B,** and **C**), 10 µm and 5 µm (**D**).

We also assessed S100A11 localization in U2OS cells, an osteosarcoma cell line featuring prominent actin SFs (Burnette et al., 2014; Hotulainen and Lappalainen 2006). Again, S100A11 localized to SFs as well FAs (Fig. 1C), although it localized primarily to peripheral adhesions at the end of dorsal SFs. Furthermore, S100A11 intensity appeared to gradually decrease with increasing distance from the FA termini of the SFs. FA localization of S100A11 was further verified by live-cell TIRF microscopy imaging of U2OS cells co-expressing GFP-S100A11 and vinculin-mCherry (Fig. 1D). Again, the TIRF images showed good overall colocalization of S100A11 and vinculin in a subset of FAs. Compared to HeLa cells, S100A11 and vinculin signals overlapped less completely. Instead, S100A11 was often enriched at the distal end of FAs or its localization extended to areas in the FA vicinity. Compared to IF images, live-cell TIRF images also showed higher levels of diffuse, non-FA-associated S100A11 across the entire basal plasma membrane, indicating that a fraction of cytoplasmic S100A11 continuously associates with the membrane and/or the cortical actin cytoskeleton network. GFP-S100A11 also localized to FA and SFs in PtK2, NIH3T3, and cancer-associated fibroblasts (CAF) cells (data not shown).

*In vitro,* S100A11 binds F-actin in a Ca^2+^-dependent manner (Sakaguchi et al., 2000; Zhao et al., 2000). To investigate if Ca^2+^-binding was also required for association with native actin SFs and FAs, U2OS cells were de-roofed using a previously established microsonication protocol (Franz and Müller, 2005). The exposed SFs (Suppl. Fig. S3) were then incubated with recombinant GFP-S100A11 protein (Suppl. Fig. S4). In presence of 1 mM Ca^2+^, GFP-S100A11 localized along SFs, while a GFP control alone only showed weak non-specific association. Importantly, when free Ca^2+^ was removed from the incubation buffer by adding 5 mM EGTA, GFP-S100A11 also displayed only weak diffuse binding, confirming Ca^2+^ dependency for SF localization. GFP S100A11-protein also displayed specific binding to exposed SFs in HeLa cells (Suppl. Fig. S5)

### Transient localization of S100A11 to disassembling FAs

Strong localization to SFs and FAs in different cell types pointed to a potential role of S100A11 in regulating these cytoskeletal structures. To obtain insight into dynamic aspects of such possible S100A11-dependent mechanisms, HeLa cells co-expressing GFP-S100A11and Lifeact-mCherry were imaged by TIRF live-cell microscopy. Only the FA-associated F-actin signal is recorded in this imaging mode, which was largely identical to experiments using *bona fide* FA markers, such as vinculin or paxillin (not shown). Timelapse movies recorded at intervals of 10 sec over a total duration of 20 min showed primarily diffuse S100A11 localization to the basal cell membrane and specific FA localization at steady intensity levels (Suppl. Movie S1). However, occasionally we also observed defined areas of transiently enhanced S100A11 intensity (Fig. 2A, Suppl. Movie S1). These striking S100A11 flashes typically occurred at or close to FAs at the cell periphery and typically lasted between 10 and 90 sec for an average time of 56 ± 31 sec (mean ± SD, n = 5). In some cases, S100A11 flashes were highly localized in small areas encompassing a single FA (Fig. 2A and 2C), while in other cases the areas of transient S100A11 recruitment were broader and covered several adjacent FAs (Fig. 2B). From the recorded timelapse movies (Suppl. Movie S1) we noticed invariable loss of FAs in the areas experiencing an S100A11 flash, either of individual FAs in small area flashes (Fig. 2A and 2C), or of several adjacent FAs in larger area flashes (Fig. 2B). To investigate the relationship between transient S100A11 recruitment and FA disassembly, kymographs were generated along trajectories of peripheral FAs targeted by S100A11 flashes (Fig. 2D and 2E and Suppl. Fig. S6). The kymographs revealed that FA disassembled within 312 ± 91 sec (mean ± SD, n = 5) after an S100A11 flash. Pre-flash FAs displayed a slow “sliding” behavior typical for FAs in stationary cells (Smilenov et al., 1999) at a speed of 0.06 ± 0.01 µm/min (mean ± SD). In contrast, after S100A11 flashing the FA translation speed increased to 0.45 ± 0.23 µm/min, or about 7-fold (Fig. 2E). A previously identified force-dependent FA disassembly mechanism involves increased actomyosin-mediated pulling forces and FAs retraction speeds (Crowley and Horwitz, 1995; Webb et al., 2004). Likewise, the elevated translocation speeds of FAs after S100A11 flashing suggested that these FAs may disassemble under increased intracellular pulling forces.

**Fig. 2.**
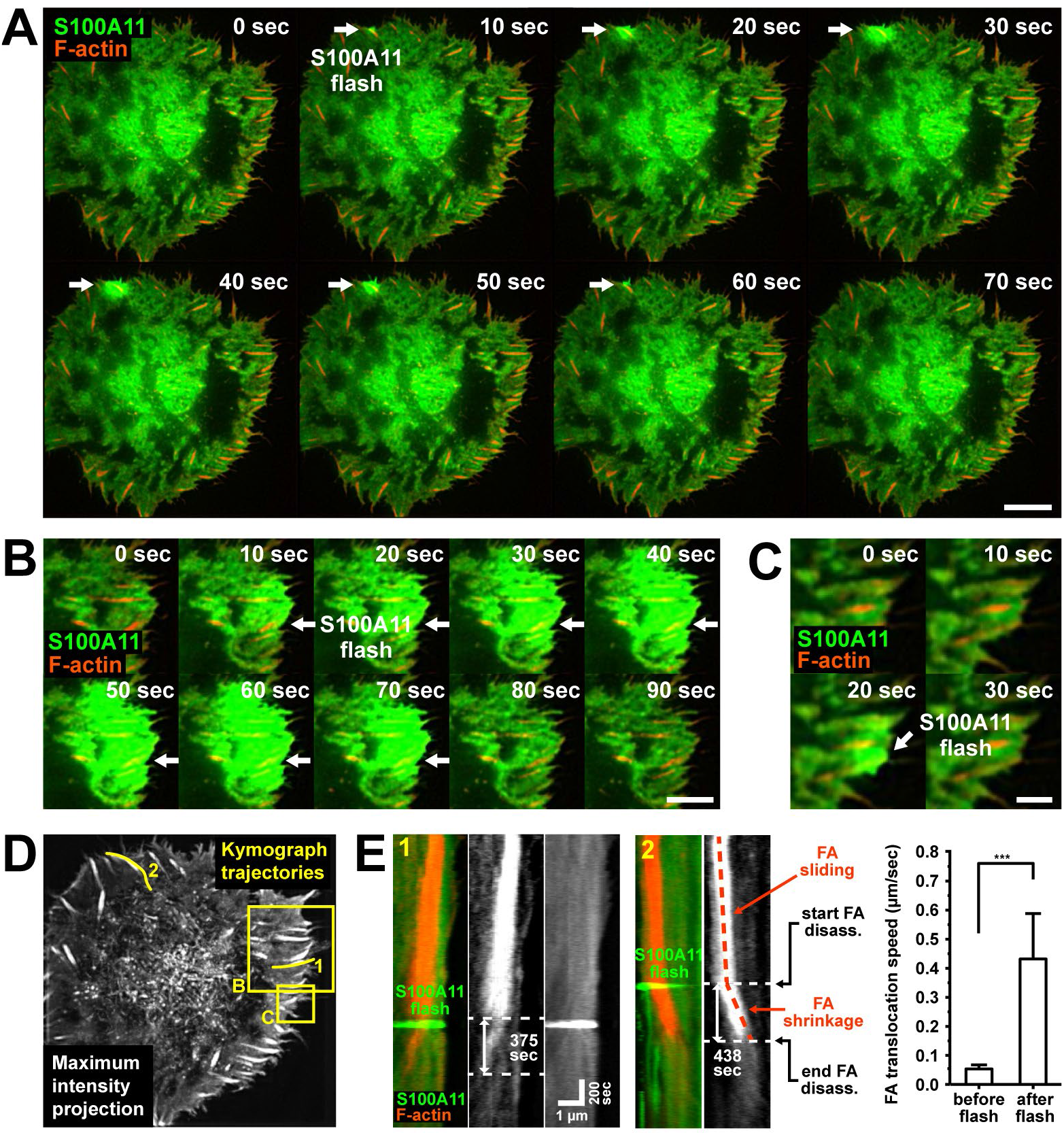
S100A11 flashes precede focal adhesion disassembly. **(A)** Selected frames from a live-cell TIRF imaging series of a single HeLa cell co-transfected with Lifeact-mCherry and GFP-S100A11. The white arrow indicates an S100A11 flash covering a single FA and lasting approx. 60 sec. Scale bar: 10 µm. **(B)** Additional frames from the same timelapse series from a cellular subregion corresponding to the yellow box B in **(D)** showing a large-area S100A11 flash covering several neighboring FAs and lasting appr. 90 sec. Scale bar: 10 µm. **(C)** Cropped region corresponding to the yellow box C in **(D)** showing a short (20 sec) localized S100A11 flash. Scale bar: 10 µm. **(D)** Maximum intensity projection of the entire timelapse stack (F-actin channel) indicating the positions of the cropped regions in **(B)** and **(C)**, as well the trajectories of two FAs used for kymograph analysis. **(E)** Two kymographs generated along the respective FA trajectories shown in **(D)**. Time values denote the time between initiation of the S100A11 flash and complete disassembly of the targeted FA. The diagram (right panel) shows the FA sliding speed before, and the FA retraction speed after the S100A11 flash (mean ± SD,) data from five S100A11 flashes.

While S100A11 flashes typically occurred over areas covering a single or several FAs, in some cases S100A11 flashes were spatially highly restricted to a small region at the distal FA end (Fig. 3A and 3B, Suppl. Movie S2). These highly localized S100A11 flashes also appeared to target specific FAs for disassembly, as a neighboring FA only several µm away displaying no flashing remained stable (Fig. 3B). Tracking the fluorescence intensities of GFP-S100A11 and vinculin-mCherry at the disassembling FA over time initially showed stable vinculin but steadily increasing S100A11 levels, followed by a sudden drop in both S100A11 and vinculin signals. These results indicated that prior to FAs disassembly S100A11 progressively accumulated at FAs until reaching a critical level, after which it was quickly released, and rapid FA disassembly commenced.

**Fig. 3.**
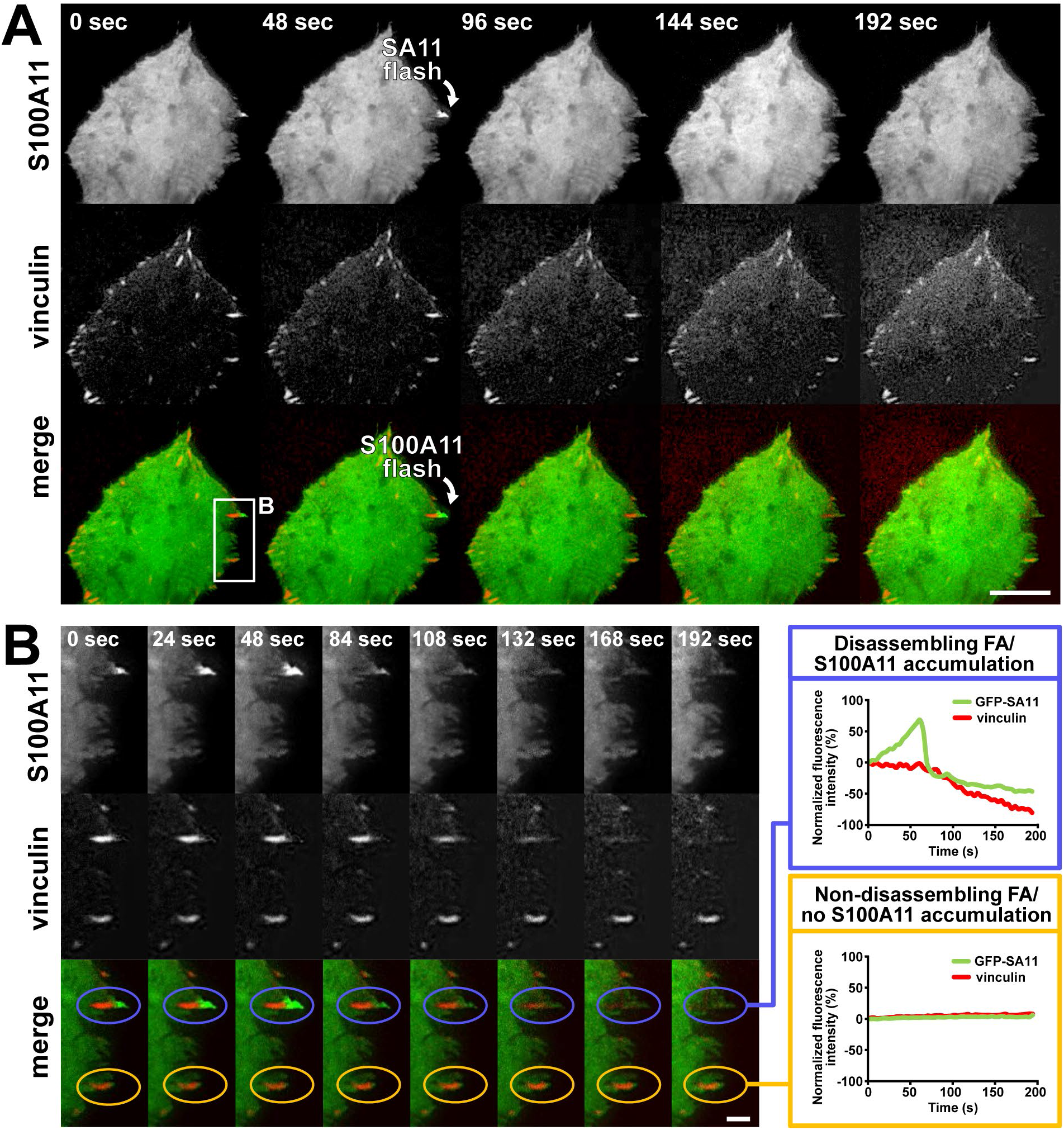
S100A11 accumulates at disassembling FAs. **(A)** Full size and **(B)** cropped frames from a live-cell TIRF imaging series of a single HeLa cell co-expressing vinculin-mCherry and GFP-S100A11. The white arrow indicates an S100A11 flash covering a single FA and lasting approx. 60 sec. Relative S100A11 and vinculin fluorescence intensities were determined in ROIs surrounding single FAs either displaying an S100A11 flash (blue ellipse) or not (yellow ellipse) and plotted over time (right panel). Scale bars: 10 µm (**A**) and 4 µm (**B**).

### Elevated intracellular Ca^2+^ recruits S100A11 to disassembling FAs

The spatial and temporal pattern of S100A11 flashing was reminiscent of intracellular Ca^2+^ flashes occurring in different cell types, (Ellefsen et al., 2019; Varadarajan et al., 2022; Wei et al., 2009; Wei et al., 2010), including during FA disassembly (Giannone et al., 2002). Ca^2+^-binding activates S100A11 by shifting the homodimer into its active, F-actin-binding conformation, and local Ca^2+^ influx may therefore be a mechanism for S100A11 recruitment to peripheral FAs. We investigated whether artificially elevating intracellular Ca^2+^ levels would stimulate S100A11 recruitment and FA disassembly. For this, we co-transfected HeLa cells with GFP-S100A11 and the fluorescent Ca^2+^-sensor R-GECO1 (Zhao et al., 2011b) for live-cell TIRF imaging. Adding the Ca^2+^ ionophore ionomycin to cells growing in Ca^2+^-containing medium strongly activated the calcium sensor within <60 sec, mirrored by a slight increase in the S100A11 signal (Fig. 4A and 4B, Suppl. Movie S3), consistent with transient Ca^2+^-dependent membrane/actin cortex recruitment of S100A11. Initial S100A11 membrane recruitment was not specifically targeted to FAs but occurred evenly across the entire membrane. However, ionomycin treatment reliably induced strong secondary, highly localized S100A11 flashes at the cell periphery within the next 5 to 10 min (Fig. 4A and 4B, Suppl. Movie S3), similar in size and duration to the random flashes observed under steady state conditions in untreated cells (Fig. 2A). During the local secondary flashes, the R-GECO1 signal also showed mild transient elevation, suggesting a Ca^2+^-dependent recruitment mechanism of S100A11 to FAs. In a variation of the previous experiments, we also incubated cells in Ca^2+^-free medium containing ionomycin. This treatment never induced S100A11 flashes, but subsequent addition of Ca^2+^ again stimulated S100A11 flashes within a similar timeframe as ionomycin addition to Ca^2+^-containing medium. Using this protocol on HeLa cells co-expressing GFP-S100A11 and vinculin-mCherry corroborated that Ca^2+^-stimulated membrane recruitment of S100A11 coincided with rapid FA translocation and FA disassembly within these areas (Fig. 4C, 4D, 4E, and Suppl. Movie S4).

**Fig. 4.**
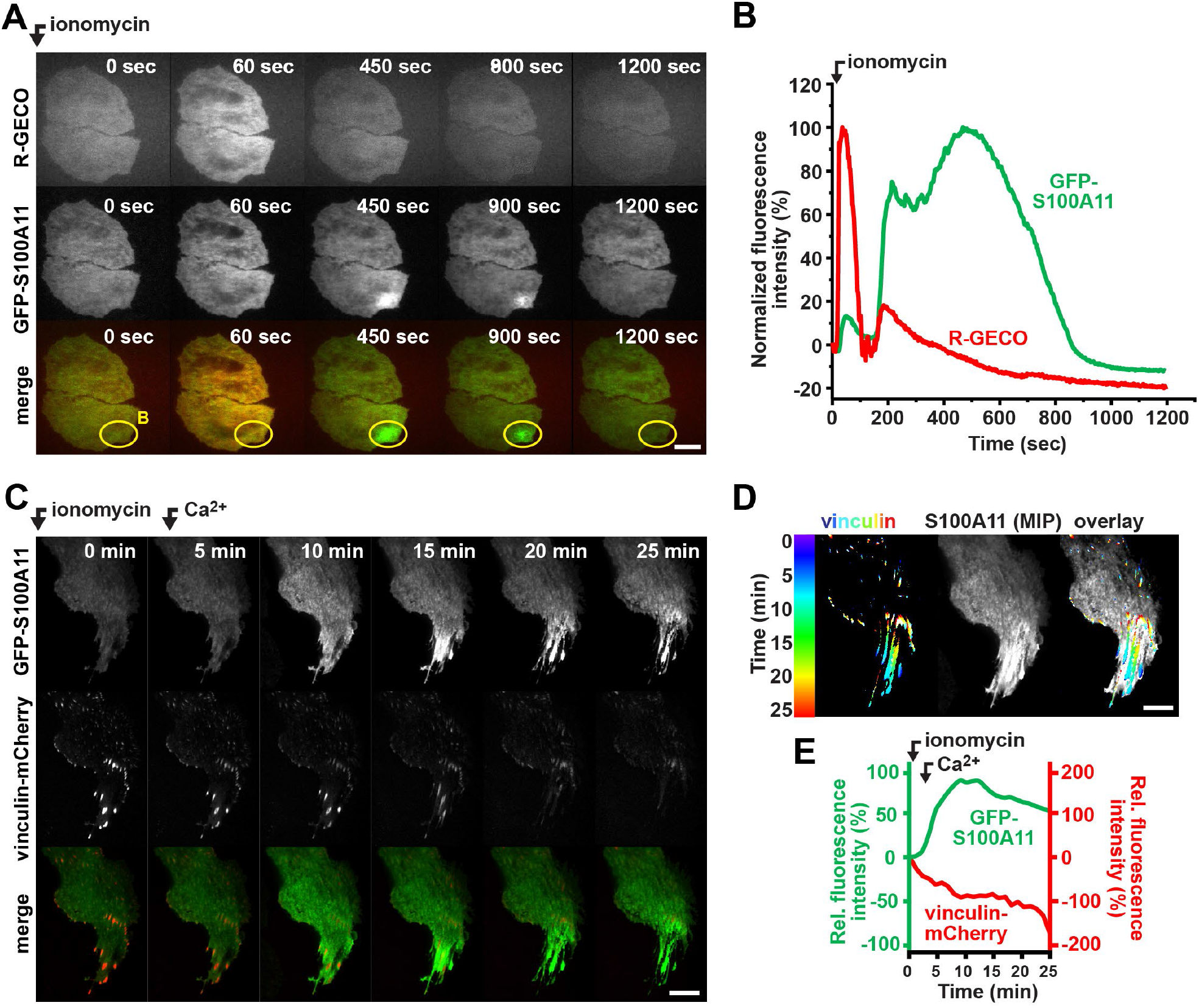
Intracellular Ca^2+^ stimulates S100A11 recruitment to disassembling focal adhesions. **(A)** Selected frames (crop) from a live-cell TIRF imaging series of HeLa cells transfected with the calcium sensor R-GECO (red) and GFP-S100A11 (green). Cells were imaged in Ca^2+^-containing medium, 3 µM ionomycin was added immediately before commencing image acquisition. Scale bar: 10 µm. **(B)** Normalized fluorescence intensity traces plotted over the full duration of the timelapse movie. **(C)** Selected frames (crop) from a live-cell TIRF imaging series of U2OS cells transfected with vinculin-mCherry (red) and GFP-S100A11 (green). Cells were imaged in Ca^2+^-free medium containing 3 µM ionomycin, and 2 mM Ca^2+^ was added 3 min after commencing image acquisition. Scale bar: 10 µm. **(D)** Maximum intensity Z-stack projection (MIP) of the complete timelapse series (S100A11 channel) overlayed with color-coded vinculin traces showing that FA translocation occurs in areas of strong S100A11 recruitment. **(E)** Relative fluorescence intensity analysis of GFP-S100A11 and vinculin-mCherry signals from the timelapse series shown in **(C)** and Suppl. Movie S4.

### Actomyosin contractility and Piezo1-mediated Ca^2+^ influx promotes S100A11 translocation to disassembling FAs

While the ionomycin experiments showed that intracellular Ca^2+^ influx stimulates S100A11 recruitment to areas of dissembling FAs, the consistent delay between initial global membrane recruitment of S100A11 immediately after ionomycin addition and the targeted localization of S100A11 to FAs suggested the requirement for an additional mechanism besides Ca^2+-^activated F-actin binding by S100A11. Specifically, the enhanced FA translocation speeds prior or during S100A11 recruitment pointed towards the possible involvement of actomyosin-driven SF contraction. In agreement, pre-incubation with the non-muscle NM II inhibitor blebbistatin fully suppressed ionomycin-stimulated S100A11 recruitment, FA translocation and FA disassembly (Fig. 5A, 5B, 5D, 5E, and Suppl. Movie S5). Elevated intracellular Ca^2+^-levels have been previously shown to stimulate actomyosin-contractility in different non-muscle cell types (Kong et al., 2019; Lee and Auersperg, 1980), likely through Ca^2+^-dependent activation of myosin light chain kinase (MLCK) (Katoh et al., 2001). Likewise, holographic tomography imaging showed that ionomycin-triggered Ca^2+^ influx strongly stimulates U2OS cell contraction and the formation of membrane blebs, cytoplasm-filled membrane extrusions driven by increased intracellular tension, while NM II inhibition by blebbistatin efficiently blocked both cell contraction and bleb formation (Suppl. Fig. S7, Suppl. Movie S7). Recent findings demonstrate that NM II-dependent traction forces transmitted by actin SFs produce highly localized Ca^2+^ spikes near tensed FAs by recruiting the force-sensitive Ca^2+^-permeable channel Piezo1 (Ellefsen et al., 2019; Yao et al., 2022). Ca^2+^ influx after contractility-dependent Piezo1 activation near tensed FA sites also provided an intriguing possible explanation for Ca^2+^-activated S100A11 recruitment to tensed FAs. Indeed, pre-treatment with the Piezo1 inhibitor GsMT4x efficiently blocked ionomycin-induced S100A11 recruitment to FAs (Fig. 5C, 5D, 5E, and Suppl. Movie S6). Our results therefore suggested that Ca^2+^ promotes S100A11 localization to FAs in two different ways: firstly, by stimulating NM II-dependent cellular contractility, leading to enhanced tensioning of FAs and local Piezo1 activation, and secondly by activating the F-actin binding capacity of S100A11 through a secondary Ca2+ signal generated by the stretch-opened Piezo1 channels.

**Fig. 5.**
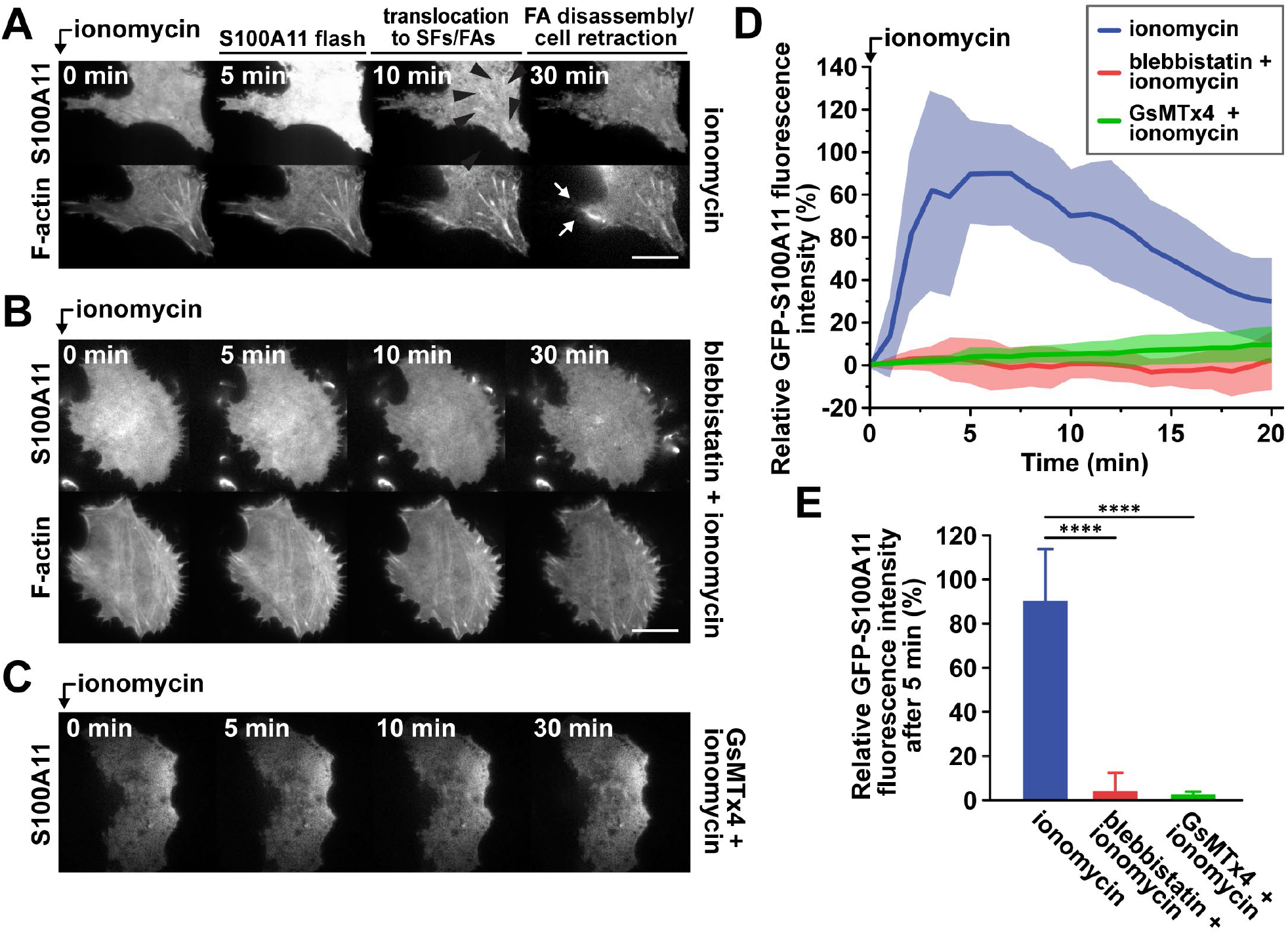
Ionomycin stimulates S100A11 recruitment to disassembling FAs by stimulating NM II-dependent cell contractility and Piezo1-mediated Ca^2+^ influx. **(A)** Selected frames from a live-cell TIRF movie of U2OS cells transfected with Lifeact-mCherry and GFP-S100A11 demonstrating S100A11 recruitment after ionomycin stimulation (3 µM) immediately before starting the image acquisition. Black arrow heads indicate S100A11 positive FA and SFs, white arrows denote areas of membrane retraction and FA translocation. **(B)** Preincubation of U2OS cells with blebbistatin (50 µM) prevents ionomycin-stimulated S100A11 recruitment, membrane retraction and FA translocation. **(C)** Preincubation with the Piezo1 inhibitor GsMTx4 (3 µM) also prevents ionomycin-stimulated S100A11 recruitment and FA translocation. **(D)** TIRF microscopy GFP-S100A11 intensity plots (mean and SD) after ionomycin stimulation after preincubation with 50 µM blebbistatin (red trace), preincubation with 3 µM GsMTx4 (green) or without preincubation (blue trace). **(E)** Relative GFP-S100A11 intensity values (mean ± SD) at 5 min after ionomycin stimulation. Data from six timelapse movies per condition.

We further investigated biomechanical aspects of S100A11 recruitment using a previously established micropipette-based pulling assay for exerting pulling forces onto individual FA sites (Riveline et al., 2001). In this assay a glass micropipette is first lowered onto the perinuclear region of a target cell, and then gently pushed against the nuclear region (Fig. 6A), thereby applying tension forces onto peripheral FAs through interconnected SFs. First, we tested whether external pulling forces would recruit S100A11 to tensed FAs. Indeed, micropipette pulling induced S100A11 accumulation to stressed FAs within a similar time (3-5 min) as Ca^2+^-stimulated SF contraction (Fig. 6B and 6C, Suppl. Movie S8 and S9). Interestingly, S100A11 again localized to the distal FA end, similar to what was observed during random S100A11 flashes in unstimulated cells (Fig. 3A and 3B). In contrast, micropipette pulling did not induce S100A11 recruitment in Ca^2+^-free medium, demonstrating a strict requirement for local S100A11 activation by Ca^2+^ (Fig. 6D). Likewise, no S100A11 recruitment occurred in Ca^2+^-containing medium in presence of the Piezo1 inhibitor GsMT4x (Fig. 6E). We furthermore tested if external pulling could bypass the requirement for NM II-driven contractility, using cells incubated with blebbistatin (Fig. 6F) or a NM IIA KO cell line (Fig. 6G). In both cases external pulling still induced S100A11 recruitment to peripheral FAs, suggesting that the required role of NM II in S100A11 recruitment is limited to the establishment of SF contractility. However, recruitment in NM II-deficient cells appeared weaker compared to control cells, raising the possibility that additional functions of NM IIA, or that other NM III isoforms also contribute to S100A11 translocation. Finally, applying external forces also recruited Piezo1-GFP to stressed FAs (Fig. 6H), suggesting that the local Ca^2+^ influx near these structures results from both stretch-dependent localization and activation of Piezo1.

**Fig. 6.**
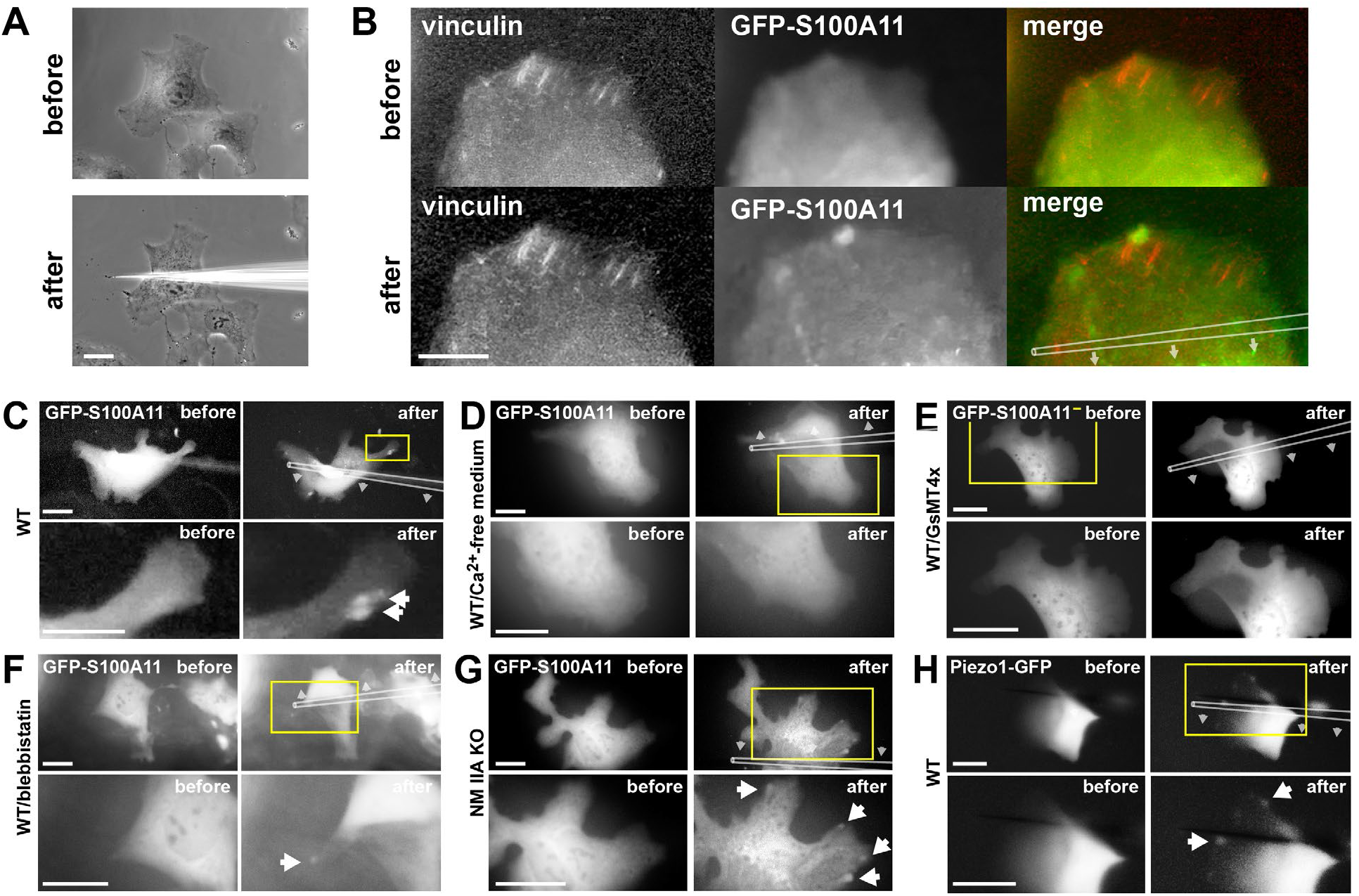
External pulling forces recruit S100A11 to focal adhesions. **(A)** Phase contrast image of U2OS cells before (upper image) and after (lower image) external pulling force application using a glass micropipette and a manual micropositioning system. **(B)** Cropped image of a single U2OS cell expressing GFP-S100A11 (green) and vinculin-mCherry (red). The original position of the micropipette before force application and the pulling direction are schematically shown. External pulling force experiments of a wildtype U2OS cell expressing GFP-S100A11 in **(C)** Ca^2+^-containing medium, **(D)** Ca^2+^-free medium, **(E)** in presence of 3 µM GsMTx4, or **(F)** in presence of 50 µM blebbistatin**. (G)** Pulling experiment of a U2OS NM IIA KO cell. **(H)** Pulling experiment in wildtype U2OS cell expressing Piezo1-GFP. Scale bars: 10 µm.

### S100A11 promotes FA disassembly during cell membrane retraction

While the previous experiments demonstrated NM IIA- and Ca^2+^-dependent recruitment of S100A11 to peripheral FAs disassembling under force, it remained unclear if S100A11 contributed to the FA disassembly process itself. To further investigate its role, we generated an S100A11 knockout U2OS cell line (S100A11 KO) and verified absence of expression by western blotting (Fig. 7A). We also analyzed FA morphology in wildtype and S100A11 KO cells (Fig. 7B-7E). Compared to wildtype cells, S100A11 KO cells displayed a larger average individual FA area (Fig. 7C), caused primarily by a subpopulation of vastly enlarged FAs (>2.5 µm^2^) absent from wildtype cells (Fig. 7D). S100A11 KO cells also featured an increased total FA area per cell (Fig. 7E). S100A11 KO cells featured a striking FA arrangement pattern characterized by dramatically elongated FAs extending along almost the entire length of thick, peripheral SFs, whereas wildtype cells rarely displayed this phenotype (Suppl. Fig. S8). Re-expression of GFP-S100A11 on the other hand fully rescued the KO phenotype and reverted FA morphology and sizes to wildtype levels (Fig. 7B-7E). FAs are perpetually remodeled, and cell type-specific FAs sizes reflect the balance between assembly and disassembly rates (Webb et al., 2002). Enlarged FAs in SA100 KO cells could result from impaired FA disassembly. U2OS are comparatively stationary cells with some stable central FAs displaying little movement over several hours, but peripheral FAs disassemble rapidly during membrane retraction (Suppl. Movie S10). Analyzing translocation velocities of peripheral FAs in retracting membrane regions in wildtype (Fig. 7F) and knockout cells (Fig. 7G) over a 60 min observation period showed significantly reduced FA retraction speeds in S100A11 KO cells (Fig. 7I, Suppl. Movie S11). Moreover, translocating FAs within membrane retractions remained large in S100A11 KO cells but shrank in wildtype cells (Fig. 7H and 7I). Reduced FA translocation and disassembly rates in S100A11 KO cells support a role for S100A11 in NM II contractility-dependent FA disassembly during membrane retraction.

**Fig. 7.**
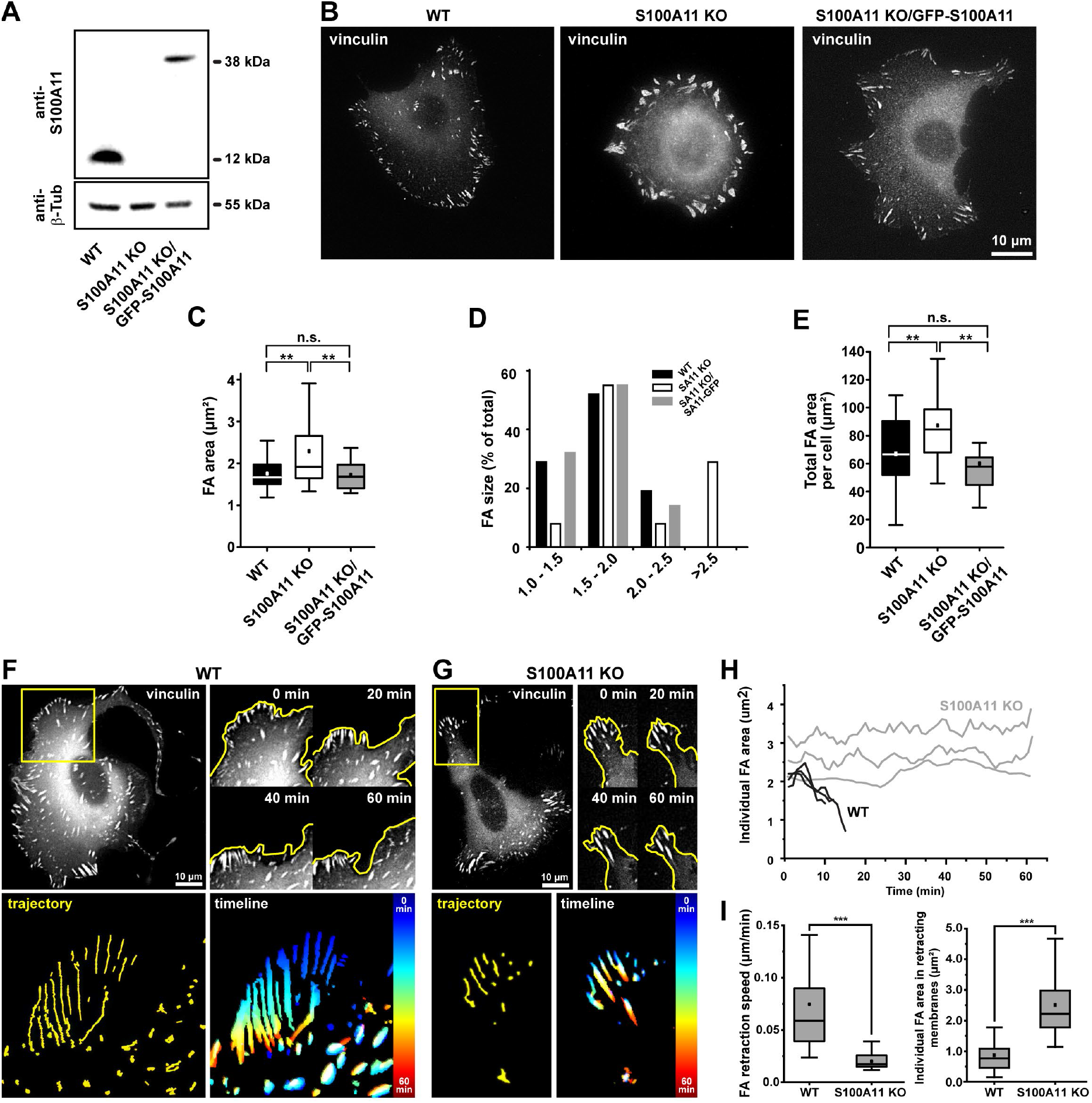
S100A11 promotes focal adhesion disassembly. **(A)** Western blot analysis of endogenous S100A11 expression in U2OS wildtype and U2OS S100A11 KO cells, as well as GFP-S100A11 rescue expression in knockout cells. The blot was re-probed against β-tubulin as a loading control. **(B)** Vinculin staining of U2OS wildtype (WT), S100A11 KO, and KO/GFP-S100A11 rescue cells. Characterization of individual FA area **(C)**, FA size distribution **(D)**, and total FA area per cell **(E)** in wildtype, KO, and rescue cells. 70 FAs from at least 20 cells per cell type analyzed. **(F)** First image frame of an epifluorescence timelapse series of a U2OS wildtype cell expressing vinculin-mCherry as a FA marker (upper left panel). Cropped still images of an area displaying active membrane retraction (yellow box) at several timepoints (upper right panel). The yellow line indicates the retracting cell border. Trajectories of translocating FAs (lower left panel) and color-coded timeline of FA displacement (lower right panel). **(G)** Corresponding images obtained for a representative S100A11-KO cells. **(H)** Size of three representative FAs (WT and KO) plotted over time. **(I)** Quantification of FA translocation speed (left box plot) and FA size (right box plot) in wildtype (WT) and SA100 KO cells.

## Discussion

Here we report a novel localization of S100A11 to FAs, both dynamically to disassembling peripheral FAs, and at comparatively steady levels at stable, stationary FAs. These two different localization patterns suggest a dual role for S100A11 in both FA maintenance and disassembly. Strikingly, S100A11 levels increased sharply at the onset of peripheral FA disassembly in a process requiring NM II-based contractility and Piezo1-dependent Ca^2+^ influx at stressed adhesion sites. Our results therefore reveal a mechano-sensitive role of S100A11 and implicate it in intracellular tension-driven FA disassembly mechanism (Fig. 8).

**Fig. 8.**
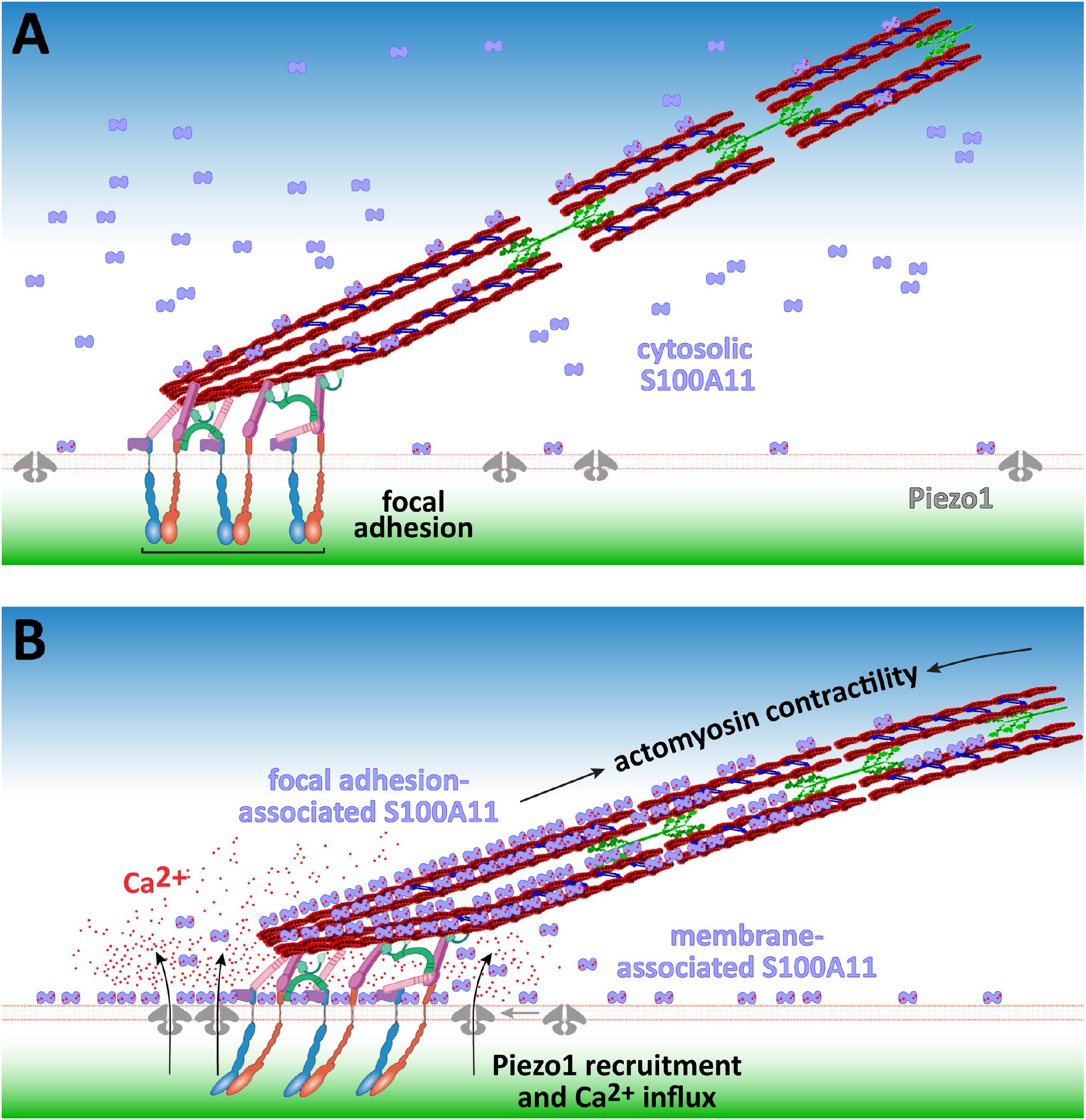
Putative model of contractility-, Piezo1- and Ca^2+^-dependent recruitment of S100A11 to FAs. **(A)** Under steady-state conditions, a small fraction of cytosolic S100A11 localizes to SFs and FAs. **(B)** NM II-mediated SF contraction increases tension at FAs and leads to the recruitment and activation of mechanosensitive Piezo1 channels. Consequently, Ca^2+^ entering through permissive Piezo1 channels activates S100A11 in the FA vicinity and promotes transient binding to F-actin structures in or near FAs, thereby promoting FA disassembly.

FAs are dynamic cell-matrix adhesion sites undergoing continuous rearrangement during cell spreading, shape changes, and migration (Webb et al., 2002), and actomyosin contractility is a known modulator of FA turnover (Aureille et al., 2023). In migrating cells, FA disassembly typically occurs at the rear, facilitating cell membrane retraction and forward propulsion (Broussard et al., 2008; Kirfel et al., 2004). Cell membrane retraction requires actomyosin contractility (Chen, 1981), and there is abundant evidence that NM II-dependent contractile forces also contribute to peripheral FA translocation and disassembly (Crowley and Horwitz, 1995; Wolfenson et al., 2011). However, the role of NM II in regulating FA lifetime is complex, since NM II activity is also required for FA growth and stabilization (Chrzanowska-Wodnicka and Burridge, 1996; Hirata et al., 2015; Oakes et al., 2012; Ridley and Hall, 1992). A resolution of this apparent discrepancy may reside in the fact the molecular composition of FAs itself is force-sensitive (Kuo et al., 2011; Wolfenson et al., 2011), and increasing NM II-mediated contraction forces above a certain threshold may shift the balance between FA stabilizers and destabilizers.

Interestingly, a previous proteomics screen identified S100A11 as part of the NM II-responsive proteome of FAs (Kuo et al., 2011). However, this study found an inverse relation between NM II-activity and S100A11 integration into FAs, in contrast to our findings. S100A11 was also detected in isolated FAs in a later proteomics study (Huang et al., 2014). The short-lived, transient recruitment regime of S100A11 suggests a function of S100A11 in initiating FA disassembly, rather than regulating the disassembly process itself. Similarly, during plasma membrane wound healing, Ca^2+^-activated S100A11 rapidly translocates to the injury site, but S100A11 levels then drop back to base levels within less than 30 s (Ashraf and Gerke, 2022), likely before completion of actin-driven membrane resealing. S100A11 may thus play functional roles in initiating dynamic actin remodeling processes in different cellular contexts.

In some cases, S100A11 localized predominantly to the distal tip of disassembling FAs. According to traction force experiments with high spatial resolution, peak traction in individual FAs is shifted towards the distal FA end (Plotnikov et al., 2012), further supporting a traction force-dependent recruitment mechanism of S100A11. Occasionally, S100A11 also localized to broader areas surrounding individual or groups of disassembling FAs, suggesting that S100A11 may also regulate cellular structures in the FA vicinity. FAs are intricately linked to surrounding cortical F-actin (Vignaud et al., 2021), and FA disassembly may require severing or remodeling such cytoskeletal crosslinks. Alternatively, membrane ripping may occur during fast FA translocation and disassembly (Kirfel et al., 2004), which could trigger the well-established role of S100A11 in plasma membrane wound healing. However, we usually observed strongest S100A11 localization signals just prior to rapid FA translocation, when membrane tears are unlikely to have occurred. In any case, localization to FAs, SFs, and membrane areas adjacent to disassembling FAs points towards a multifunction role of S100A11 in FA regulation.

While transient S100A11 localization to FAs reliably predicted subsequent FAs disassembly, and aberrant FA morphology and delayed FA disassembly during membrane retraction in S100A11 KO cells confirmed a functional role of S100A11 in FA disassembly, the underlying molecular mechanisms, as well as its binding partners in and around the FA plaque, remain to be elucidated. Surprisingly, global Ca^2+^ influx after ionomycin treatment initially provoked only weak, diffuse S100A11 membrane recruitment, while specific recruitment to FA sites required a secondary Ca^2+^ signal through tension-activated Piezo1 channels in the FA vicinity. Apparently, S100A11 recruitment to specific cellular compartments requires precise local and temporal control of free Ca^2+^-levels. Along similar lines, local rather than global Ca^2+^ increases are required for FA disassembly (Giannone et al., 2004). Successful isolation of S100A11-annexin A1 complexes also requires precisely controlled cellular Ca^2+^ thresholds (Gerke and Moss, 2002), further underlining the Ca^2+^-concentration dependency of S100A11 interactions with binding partners.

Besides Ca^2+^-activated F-actin binding, additional mechano-regulated mechanisms may target S100A11 specifically to tensed FAs. FAs contain a variety of tension-sensitive proteins and force-induced unfolding of these proteins could expose specific binding sites for S100A11 in tensed FAs. S100A11 may also serve as a docking site for the subsequent recruitment of additional FA regulators, or it may co-translocate with other proteins. Here potential candidates are annexins, which cooperate with S100A11 in the context of plasma membrane wound healing (Jaiswal et al., 2014), membrane organization (Chang et al., 2007), or endocytosis (Seemann et al., 1997).

Additional FA disassembly mechanisms have been outlined in detail, with microtubule-dependent processes receiving particular attention. Disrupting microtubules buy nocodazole reinforces SFs and stabilizes FAs (Bershadsky et al., 1996; Danowski, 1989; Liu et al., 1998), while microtubule repolymerization after nocodazole wash-out quickly induces global FA disassembly (Ezratty et al., 2005). Likewise, repeated microtubule contact targets specific FAs for disassembly (Kaverina et al., 1998; Small et al., 2002; Small and Kaverina, 2003), for instance by stimulating focal adhesion kinase, dynamin-, and clathrin-mediated endocytosis of FA components (Chao and Kunz, 2009; Ezratty et al., 2009). Moreover, Ca^2+^ influx-dependent mechanisms have been identified, in particular calpain-mediated cleavage of FA components (Huttenlocher et al., 1997), which however can also act downstream of microtubule-induced FA disassembly pathways (Bhatt et al., 2002). Similar to Ca^2+^-mediated FA disassembly after ionomycin-stimulation, S100A11 also localizes to disassembling FAs following nocodozole washout (our observation), suggesting that both pathways share common molecular mechanisms involving contractility regulation and local Ca^2+^ influx. In agreement, recent findings have established a mechanistic link between microtubule- and NM II-dependent FA disassembly (Aureille et al., 2023).

Several recent studies have identified tension-dependent recruitment and activation of Piezo1 channels to FAs (Ellefsen et al., 2019) and implicated the resulting local Ca^2+^ signals in the modulation of FA function (Yao et al., 2022). In agreement, our micropipette assay also demonstrates Piezo1 recruitment to tensed FAs in response to an external force. Piezo1 activation at tensed FAs may either result from enhanced membrane tension, or via direct linkage of Peizo1 to still unknown FA components (Yao et al., 2022). Asymmetric local Ca^2+^ flickers have previously been identified as a mechanism to steer cell migration (Wei et al., 2009), likely because they target specific FAs for disassembly. MLCK-dependent phosphorylation of myosin II has been identified as the force-producing mechanism driving Piezo1 activation (Ellefsen et al., 2019). Since MLCK itself is activated by intracellular Ca^2+^, this raises the intriguing possibility of a feedback loop in which Piezo1-mediated Ca^2+^-dependent influx near tensed FAs further stimulates actomyosin contractility, leading to further Piezo activation and so on, as has been previously suggested (Ellefsen et al., 2019; Hirata et al., 2015).

Cancer cell migration and invasion crucially depend on actin cytoskeleton remodeling and adhesion modulation and actin-binding proteins are frequently dysregulated in cancer and contribute to malignancy and unfavorable clinical prognosis (Izdebska et al., 2020; Suresh and Diaz, 2021). S100A11 also has a well-established role in cancer (Bresnick et al., 2015), promoting aggressive traits such as increased pseudopodal protrusion, migration, invasion, and metastasis (Niu et al., 2016; Shankar et al., 2010). In this context, the newly identified role of S100A11 in regulating spatially controlled FA disassembly could contribute to its pro-migratory and -invasive properties. In conclusion, our study identifies a novel NM II-, Piezo1-, and Ca^2+^-dependent role for S100A11 during force-dependent FA disassembly. The novel identification of S100A11 as a mechano-responsive protein thus further expands its diverse functional repertoire.

## Abbreviations

SF: Stress fibers
FA: focal adhesions
NM II: non-muscle myosin II
MLCK: myosin light chain kinase

## Acknowledgements and Funding

CMF received support from the Japanese Ministry of Education, Culture, Sports, Science and Technology (World Premier International Research Center Initiative WPI) and through JSPS KAKENHI 20H03218. YRL received support from the Japan-Taiwan Exchange Association. YM is supported by the Cannon Foundation and the Ohsumi Frontier Science Foundation. AT received support through JSPS KAKENHI 23H02123 and 21KK0126. MB thanks the Deutsche Forschungsgemeinschaft (DFG, German Research Foundation) under Germany’s Excellence Strategy via the Excellence Cluster “3D Matter Made to Order”, EXC-2082/1-390761711. We furthermore thank Hiroki Konno, Satoshi Arai and Cong Vu (NanoLSI, Kanazawa University) for generous help with laboratory equipment and reagents.

## Supplemental Material

### Supplementary Figures

**Suppl. Fig. S1.**
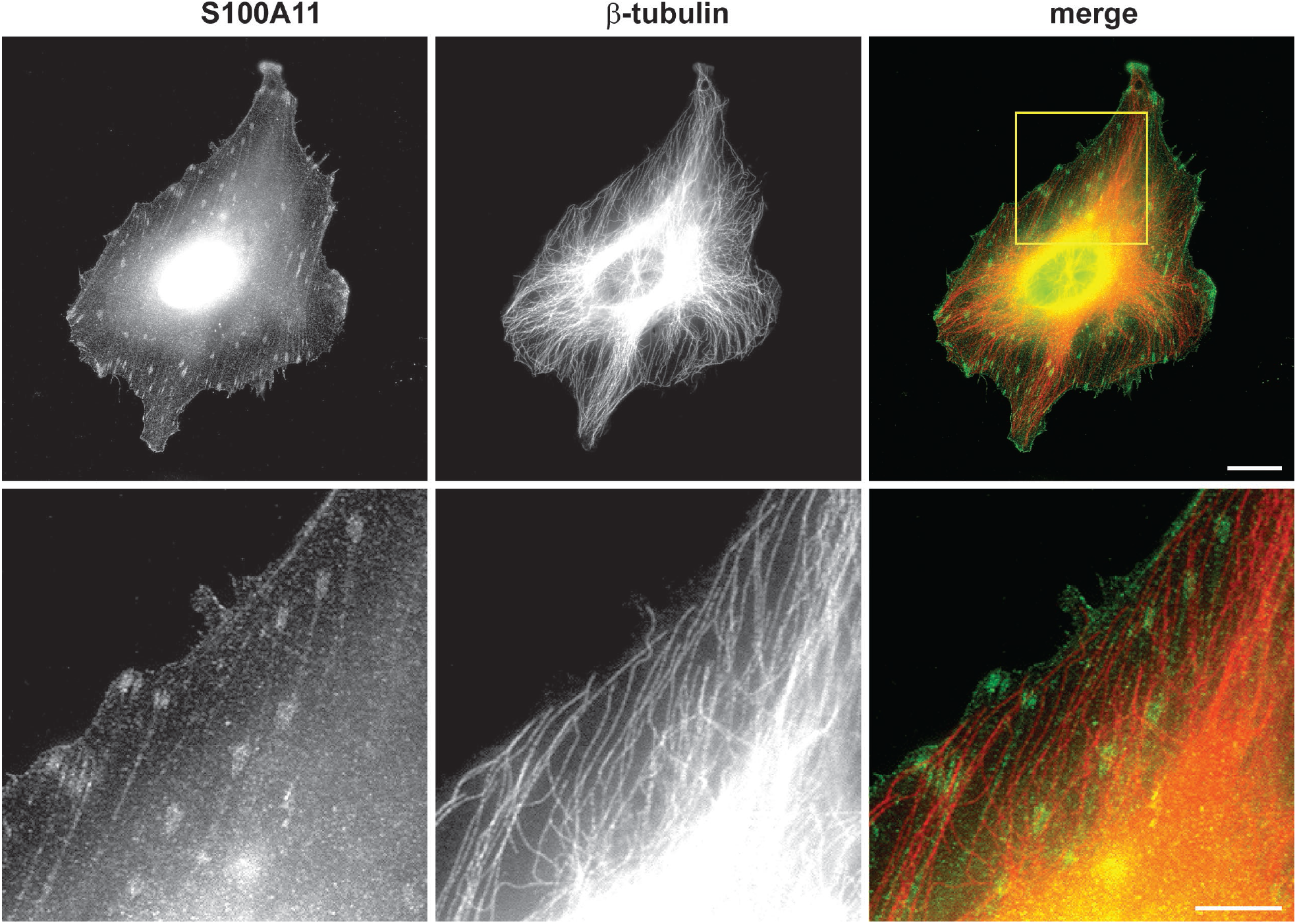
Co-immunofluorescence staining for microtubules (β-tubulin, red) and S100A11 (green) in a HeLa cell. The yellow box indicates the enlarged region displayed in the lower panel. Scale bars 10 µm (upper panel) and 5 µm (lower panel).

**Suppl. Fig. S2.**
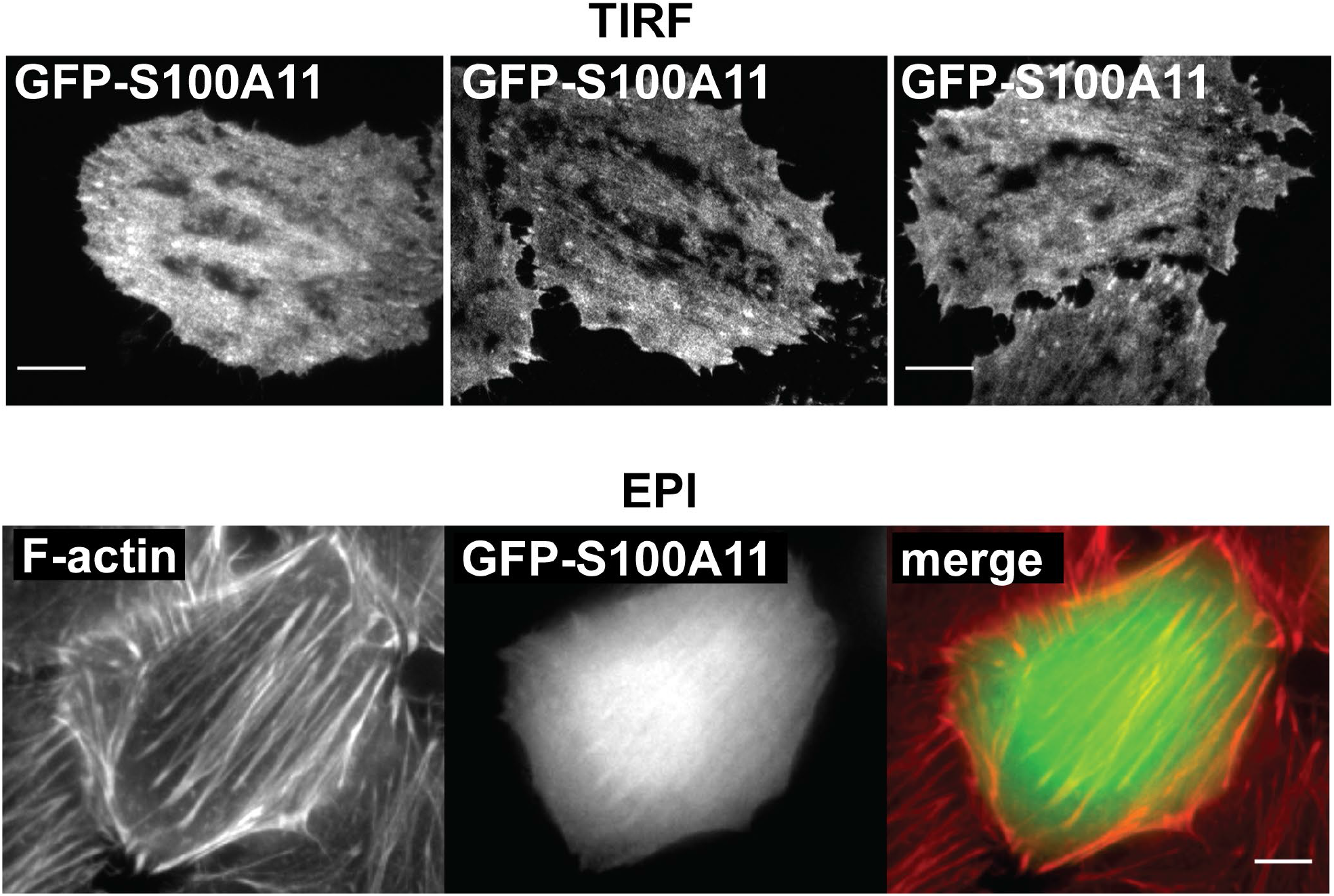
S100A11 localizes to SFs and FAs. **(Upper row)** HeLa cells were transfected with GFP-S100A11 and imaged by TIRF microscopy; three representative images are shown. **(Lower row)** HeLa cells were transfected with GFP-S100A11 and stained with phalloidin to visualize F-actin by epi-fluorescence. Scale bars 10 µm.

**Suppl. Fig. S3.**
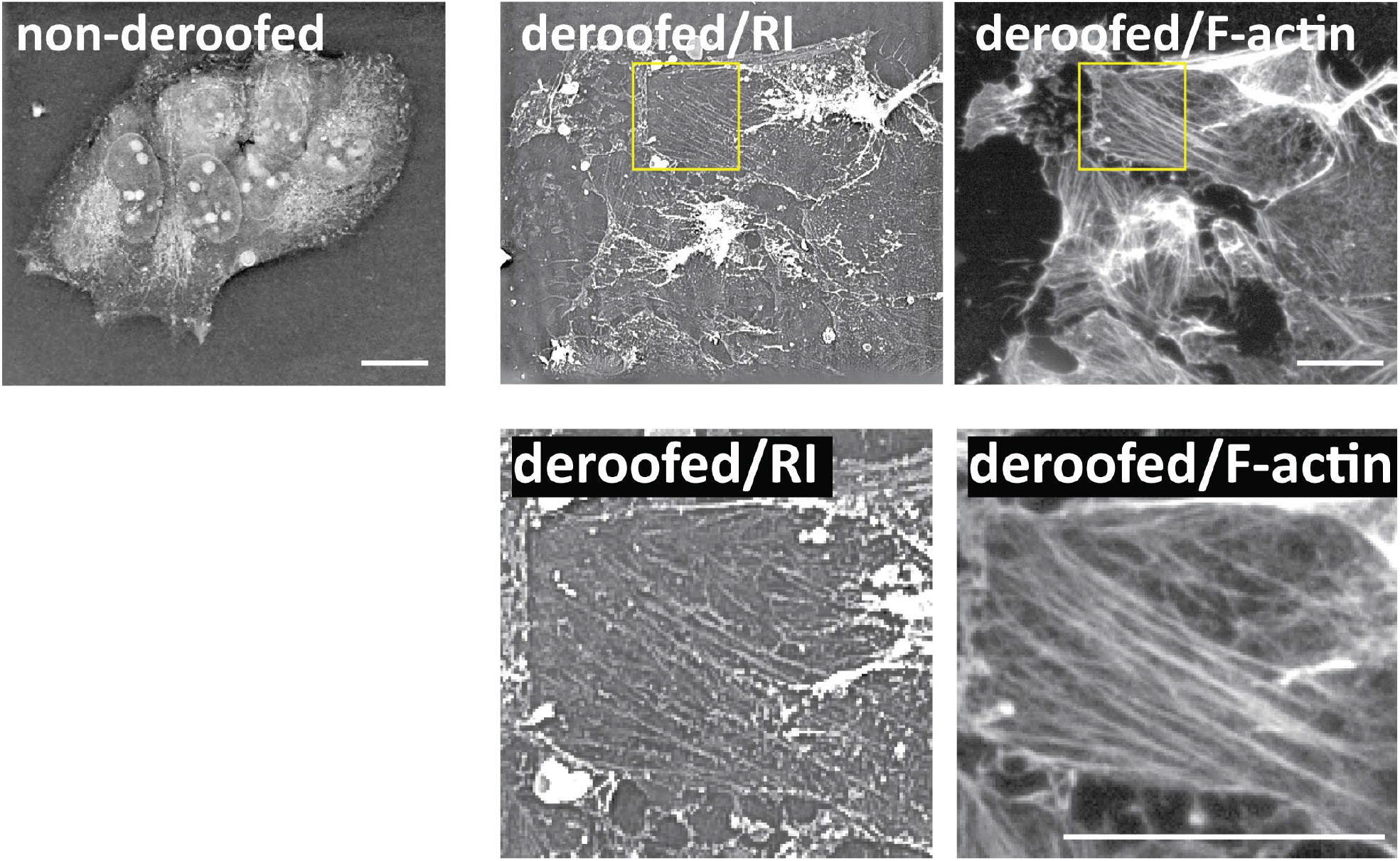
Exposing SFs in deroofed U2OS cells. Holographic tomography images (RI, Nanolive 3D Explorer) of non-deroofed (upper left panel) and de-roofed U2OS cells (upper middle panel). SFs were exposed by mechanically deroofing U2OS cells using a previous established protocol (Franz and Müller, 2005). After de-roofing cells actin SFs were stained using phalloidin-AF568 (upper right panel). The areas indicated by the yellow boxes containing well-preserved SFs are shown at increased magnification in the lower panels (left: RI, right: fluorescence). Scale bars 20 µm.

**Suppl. Fig. S4.**
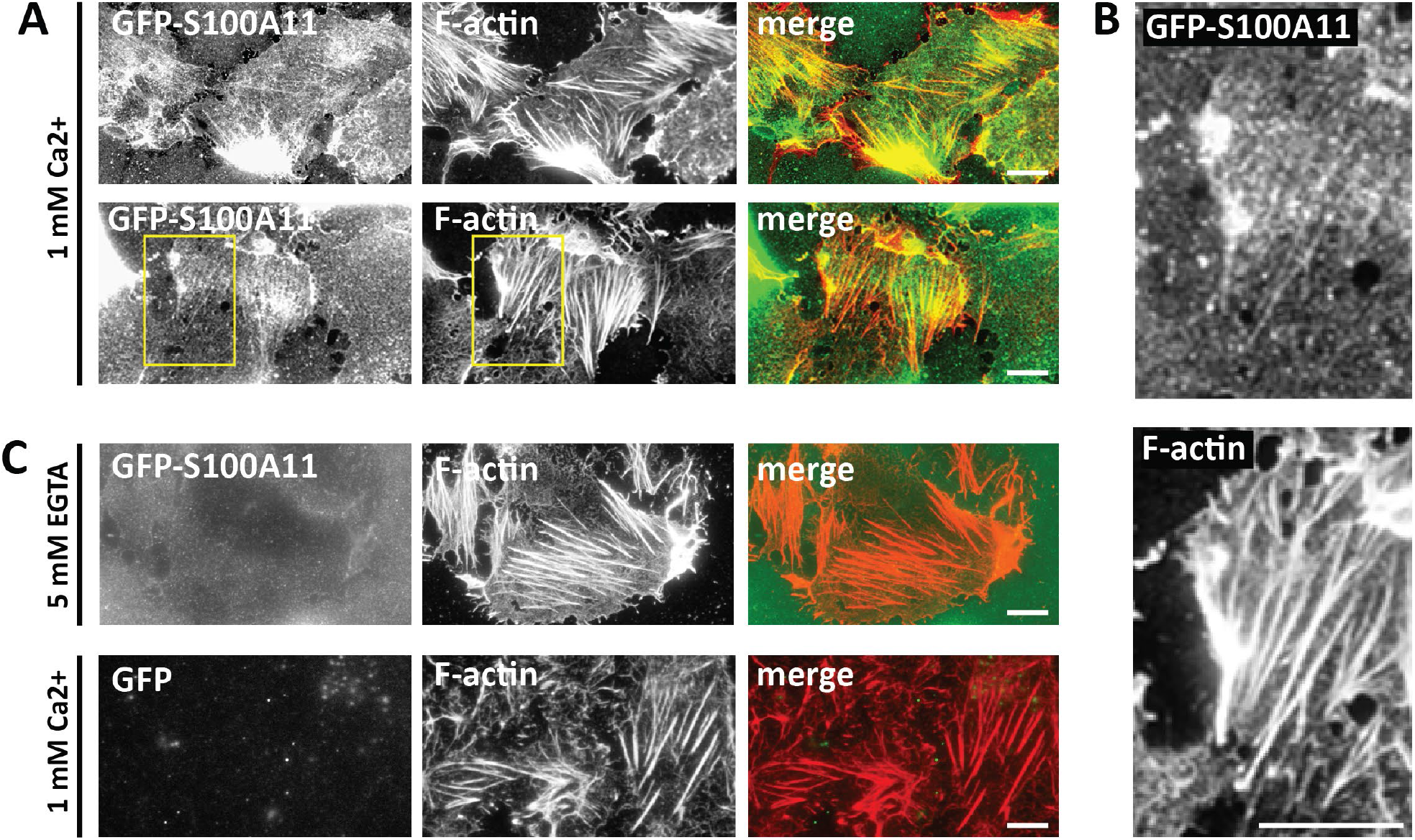
Ca^2+^-dependent interaction of S100A11 with SFs in deroofed U2OS cells. **(A)** De-roofed U2OS cell samples displaying exposed SFs incubated with purified GFP-S100A11 protein (1 µM) for 1 h in Ca2^+^-containing buffer (100 mM KCl, 2 mM MgCl_2_, 1 mM CaCl_2_, and 10 mM HEPES, pH 7.4). After 3 washes in protein-free buffer, cells were imaged with the 60x objective of an inverted fluorescence microscope (Keyence BZ-X810, two representative cell samples). **(B)** Magnified views, corresponding to the yellow boxes in **(A)**, of the GFP-S100A11 (upper panel) and F-actin (lower panel) images. **(C)** No SF association was detectable if free Ca^2+^ was removed from the incubation buffer by adding 5 mM EGTA (upper row). Furthermore, incubation with GFP protein also did not decorate SFs, demonstrating the specificity of the S100A11/F-actin interaction (lower row). Scale bars 10 µm.

**Suppl. Fig. S5.**
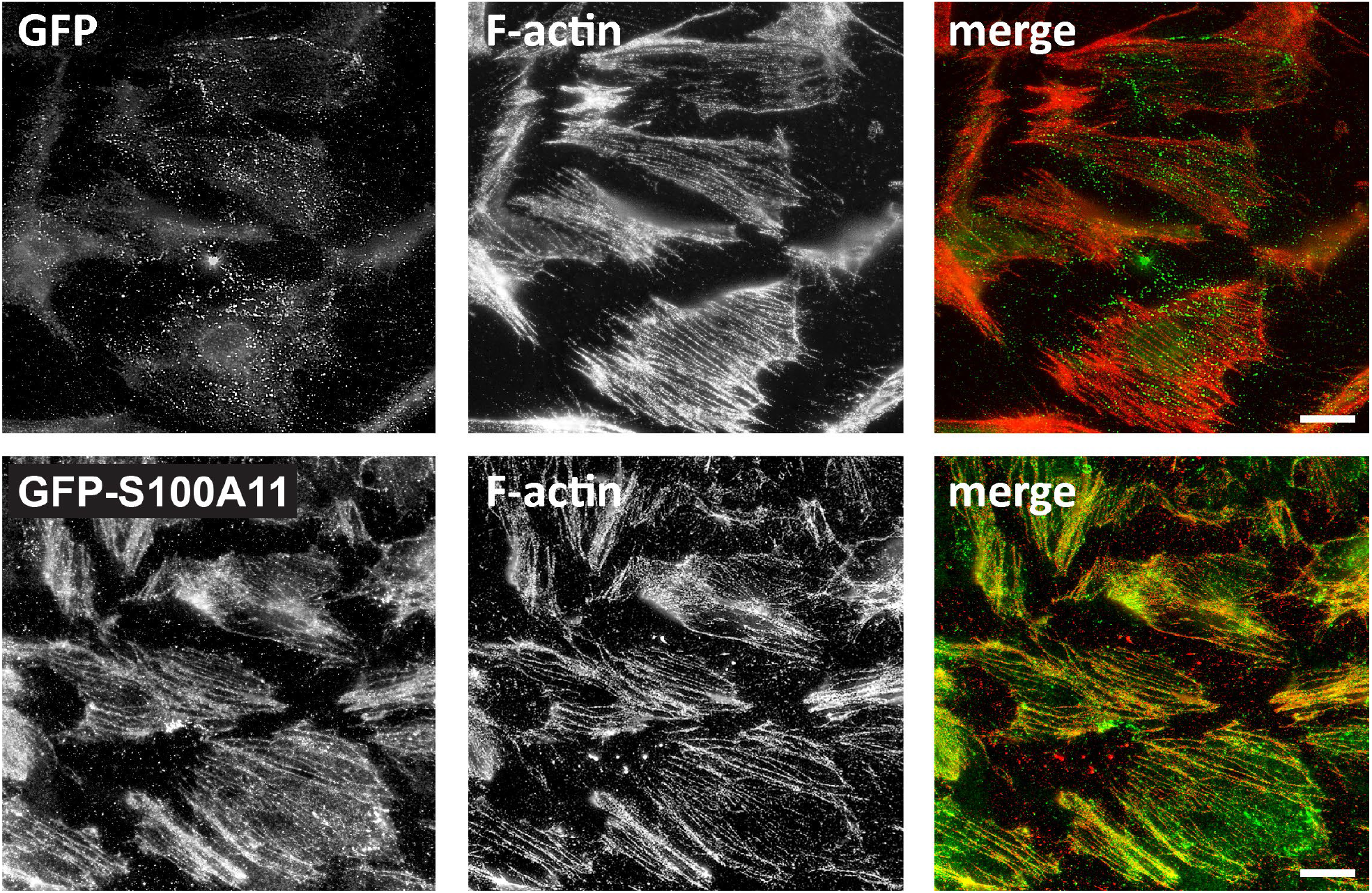
Ca^2+^-dependent interaction of S100A11 with SFs in deroofed Hela cells. De- roofed Hela cells were incubated with purified GFP (1 µM, upper row) or GFP-S100A11 protein (1 µM, lower row) for 1 h in Ca2^+^-containing buffer (100 mM KCl, 2 mM MgCl_2_, 1 mM CaCl_2_, and 10 mM HEPES, pH 7.4). After 3 washes in protein-free buffer, cells were imaged with the 60x objective of an inverted fluorescence microscope (Keyence BZ-X810, two representative cell samples).

**Suppl. Fig. S6.**
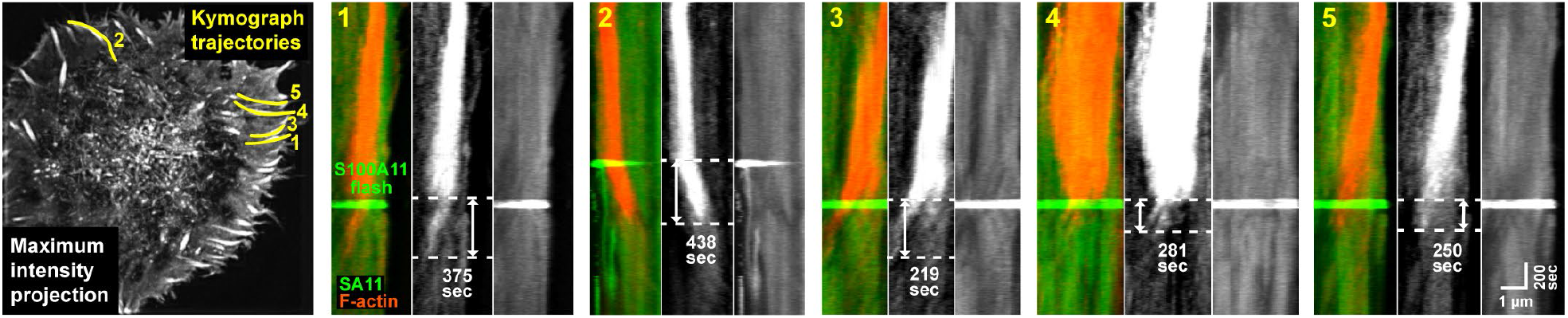
Rapid S100A11 flashes occur at sites of FA disassembly. **(A)** Maximum intensity projection of a TIRF timelapse series (F-actin channel, left most panel). The yellow lines indicate five trajectories along which kymographs were compiled. The time intervals between appearance of a GFP-S100A11 flash (green) and complete disassembly of the targeted FA (red) are indicated in the individual kymograph plots.

**Suppl. Fig. S7.**
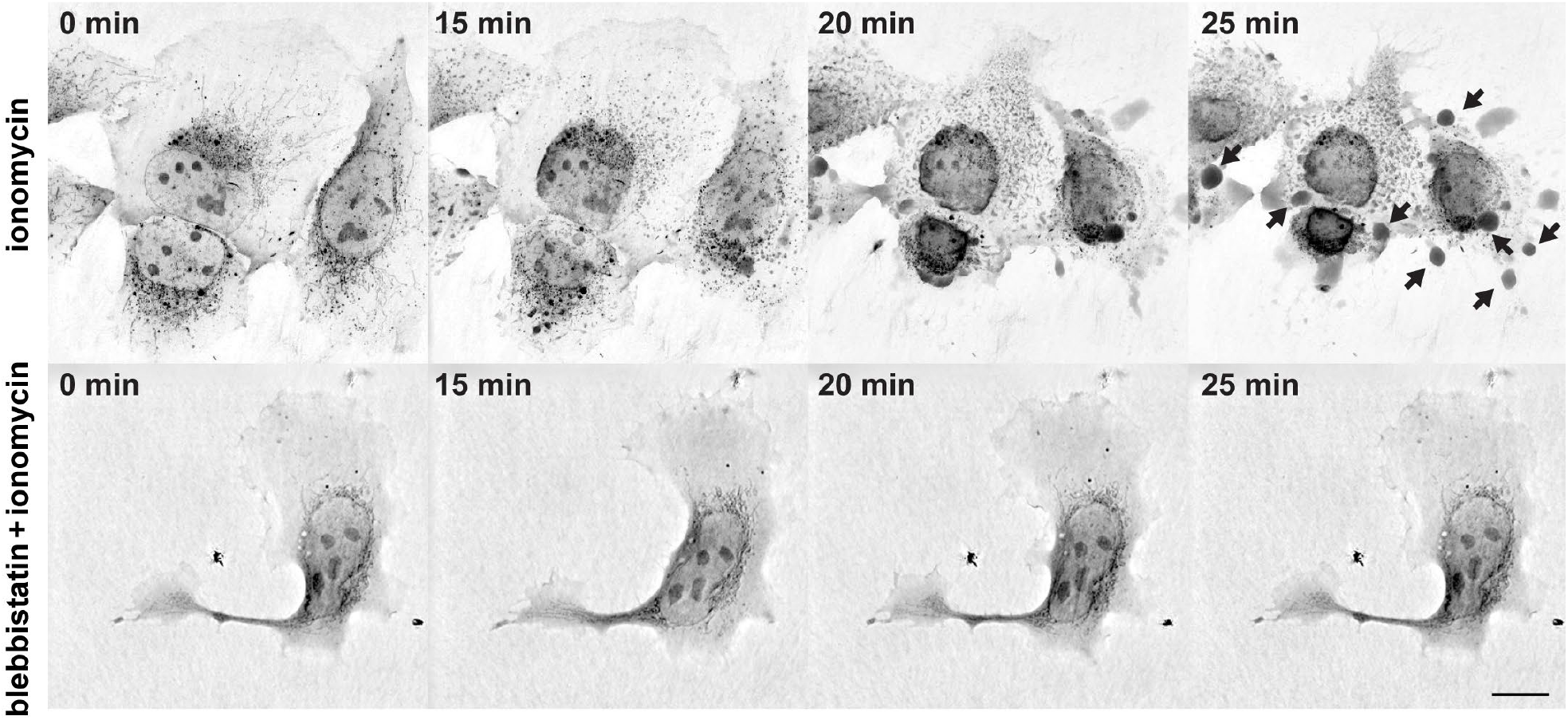
Stimulation of actomyosin-based intracellular contractility by ionomycin. **(A)** Holographic tomography timelapse recordings of U2OS cells stimulated with ionomycin (3 µM) without (top row) or with (bottom row) preincubation with 50 µM blebbistatin. Black arrows indicate membrane blebs. Scale bar: 10 µm.

**Suppl. Fig. S8.**
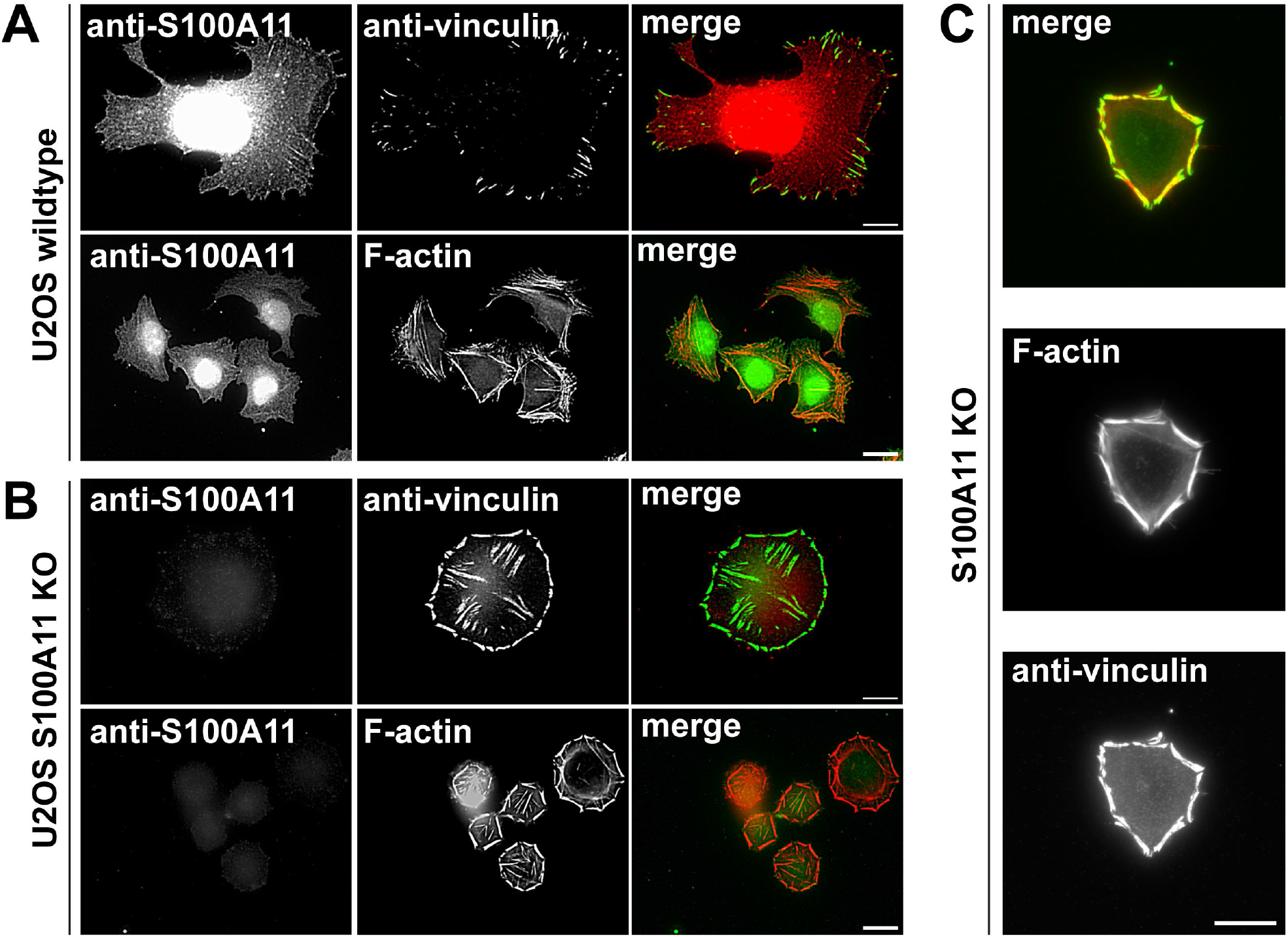
Altered SF and FA morphology in S100A11 knockout U2OS cells. **(A)** Immunofluorescence staining of U2OS wildtype cells for endogenous S100A11 and vinculin (top row), or S100A11 and phalloidin staining (bottom row). **(B)** Staining for S100A11 and vinculin (top row), or S100A11 and phalloidin staining (bottom row) in S100A11 knockout cells. **(C)** After 4 hours of spreading on fibronectin, S100A11 KO cells frequently display an unusual morphology characterized by strongly enlarged circumferential FA and SFs.

### Supplementary Movie Legends

**Supplementary Movie S1. S100A11 flashes at FA disassembly sites.** TIRF timelapse image series of a HeLa cell co-expressing GFP-S100A11 (green) and Lifeact mCherry (red) collected at a rate of 1 frame every 10 s. Scale Bar: 10 µm.

**Supplementary Movie S2. Spatially controlled S100A11 flashes are specific to disassembling FAs.** Uncropped (top row) and cropped (lower row) TIRF timelapse image series of a HeLa cell co-expressing GFP-S100A11 (green) and vinculin-mCherry (red) collected at a rate of 1 frame every 4 s. Movie playback rate: 5 fps; Scale Bar: 10 µm. A distinct S100A11 flash occurs distally to a FA that subsequently disassembles (blue arrow). In contrast, a neighboring FA located about 14 µm away experiences no S100A11 flash and remains stable (yellow arrow).

**Supplementary Movie S3. Ionomycin treatment stimulates Ca^2+^-mediated S100A11 recruitment to discrete regions at the cell periphery.** HeLa cells were co-transfected with GFP-S100A11 (green) and the calcium sensor R-GECO1 (red). After stimulation with 3 µM ionomycin in Ca^2+^-containing medium, a single cell was then immediately imaged at a rate of 1 frame every 20 s by TIRF microscopy. Scale bar: 10 µm.

**Supplementary Movie S4. S100A11 translocates to disassembling FA after ionomycin stimulation.** U2OS were cells co-transfected with GFP-S100A11 (green) and vinculin-mCherry (red) and imaged in Ca^2+^-free medium containing 3 µM ionomycin by TIRF microscopy at a rate of 1 frame every 5 s. 3 min after starting the image recording 2 mM Ca^2+^ was added. Scale bar: 10 µm.

**Supplementary Movie S5. Ionomycin-induced S100A11 recruitment to disassembling FAs requires NM II activity. Upper panel**: U2OS cells co-transfected with GFP-S100A11 (green) and Lifeact-mCherry (red) were stimulated with 3 µM ionomycin immediately before TIRF imaging. S100A11 displays recruitment to FAs. **Lower panel:** Presence of the NM II inhibitor blebbistatin (50 µM) inhibits ionomycin-induced GFP-S100A11 recruitment.

**Supplementary Movie S6. Ionomycin-induced S100A11 recruitment to FAs requires active Piezo1 channels.** U2OS cells transfected with GFP-S100A11 were incubated without (left) or with (right) the Piezo1 inhibitor GsMTx4 (0.06 µg/ml). Subsequent stimulation with ionomycin (3 µM) induced S100A11 recruitment only when Piezo1 channels were not inhibited.

**Supplementary Movie S7. Ionomycin induced cell contraction is blocked by NM II inhibition.** U2OS cells stimulated with 3 µM ionomycin either without (left) or with (right) blebbistatin preincubation (50 µM) were imaged by holographic tomography. Without blebbistatin, ionomycin induces cell contraction and membrane bleb formation.

**Supplementary Movie S8. External forces recruit S100A11 to stressed FAs.** Using a manual micropipette manipulation setup, external pulling forces were applied onto peripheral FAs in a U2OS cells co-transfected with GFP-S100A11 and vinculin-mCherry. The first image is presented as a merged image (green: GFP-S100A11, red: vinculin-mCherry). After the timelapse series, the first and final frames are again presented as merged imaged.

**Supplementary Movie 9. External forces induce GFP-S100A11 accumulation at stressed FAs.** Pulling forces are applied to a single U2OS cell expressing GFP-S100A11 via a glass micropipette.

**Supplementary Movie 10. Reduced FA dynamics in U2OS S100A11 KO cells.** U2OS wildtype and S100A11 KO cells were transfected with vinculin mCherry as a FA marker and imaged by epifluorescence microscopy for 1 hour.

**Supplementary Movie 11. Reduced FA translocation and disassembly within retracting membranes.** WT U2OS cells (WT, upper row) and S100A11 KO U2OS cells (SA11, lower row) transfected with vinculin-mCherry to mark FAs were imaged by timelapse epifluorescence microscopy for 1 h. The images show 40 x 40 µm^2^ subregions displaying active membrane retraction. Overlays (right images) display tracked FAs (red) and FA trajectories (yellow).

